# Enhanced mucosal mitochondrial function corrects dysbiosis and OXPHOS metabolism in IBD

**DOI:** 10.1101/2024.03.14.584471

**Authors:** Neeraj Kapur, M. Ashfaqul Alam, Syed Adeel Hassan, Parth H Patel, Lesley A. Wempe, Sarayu Bhogoju, Tatiana Goretsky, Jong Hyun Kim, Jeremy Herzog, Yong Ge, Samuel G. Awuah, Mariana Byndloss, Andreas J Baumler, Mansour M. Zadeh, R. Balfour Sartor, Terrence Barrett

## Abstract

**Background:** Mitochondrial (Mito) dysfunction in IBD reduces mucosal O2 consumption and increases O2 delivery to the microbiome. Increased enteric O2 promotes blooms of facultative anaerobes (eg. *Proteobacteria*) and restricts obligate anaerobes (eg. *Firmicutes*). Dysbiotic metabolites negatively affect host metabolism and immunity. Our novel compound (AuPhos) upregulates intestinal epithelial cell (IEC) mito function, attenuates colitis and corrects dysbiosis in humanized *Il10-/-* mice. We posit that AuPhos corrects IBD-associated dysbiotic metabolism.

**Methods:** Primary effect of AuPhos on mucosal Mito respiration and healing process was studied in ex vivo treated human colonic biopsies and piroxicam-accelerated (Px) *Il10-/-* mice. Secondary effect on microbiome was tested in DSS-colitis WT B6 and germ-free 129.SvEv WT or *Il10-/-* mice reconstituted with human IBD stool (Hu-*Il10-/-*). Mice were treated orally with AuPhos (10- or 25- mg/kg; q3d) or vehicle, stool samples collected for fecal lipocalin-2 (f-LCN2) assay and microbiome analyses using 16S rRNA sequencing. AuPhos effect on microbial metabolites was determined using untargeted global metabolomics. AuPhos-induced hypoxia in IECs was assessed by Hypoxyprobe-1 staining in sections from pimonidazole HCl-infused DSS-mice. Effect of AuPhos on enteric oxygenation was assessed by *E. coli Nissle 1917 WT* (aerobic respiration-proficient) and *cytochrome oxidase (cydA)* mutant (aerobic respiration-deficient).

**Results:** Metagenomic (16S) analysis revealed AuPhos reduced relative abundances of *Proteobacteria* and increased blooms of *Firmicutes* in uninflamed B6 WT, DSS-colitis, Hu-WT and Hu-*Il10-/-* mice. AuPhos also increased hypoxyprobe-1 staining in surface IECs suggesting enhanced O2 utilization. AuPhos-induced anaerobiosis was confirmed by a significant increase in cydA mutant compared to WT (O2-utlizing) *E.coli*. Ex vivo treatment of human biopsies with AuPhos showed significant increase in Mito mass, and complexes I and IV. Further, gene expression analysis of AuPhos-treated biopsies showed increase in stem cell markers (Lgr4, Lgr5, Lrig1), with concomitant decreases in pro-inflammatory markers (IL1β,MCP1, RankL). Histological investigation of AuPhos-fed Px-*Il10-/-* mice showed significantly decreased colitis score in AuPhos-treated Px-*Il10-/-* mice, with decrease in mRNA of pro-inflammatory cytokines and increase in Mito complexes (*ND5*, *ATP6*). AuPhos significantly altered microbial metabolites associated with SCFA synthesis, FAO, TCA cycle, tryptophan and polyamine biosynthesis pathways. AuPhos increased pyruvate, 4-hydroxybutyrate, 2-hydroxyglutarate and succinate, suggesting an upregulation of pyruvate and glutarate pathways of butyrate production. AuPhos reduced IBD-associated primary bile acids (BA) with concomitant increase in secondary BA (SBA). AuPhos treatment significantly decreased acylcarnitines and increased L-carnitine reflective of enhanced FAO. AuPhos increases TCA cycle intermediates and creatine, energy reservoir substrates indicating enhanced OxPHOS. Besides, AuPhos also upregulates tryptophan metabolism, decreases Kynurenine and its derivatives, and increases polyamine biosynthesis pathway (Putresceine and Spermine).

**Conclusion:** These findings indicate that AuPhos-enhanced IEC mitochondrial function reduces enteric O2 delivery, which corrects disease-associated metabolomics by restoring short-chain fatty acids, SBA, AA and IEC energy metabolism.

**Graphical abstract:** 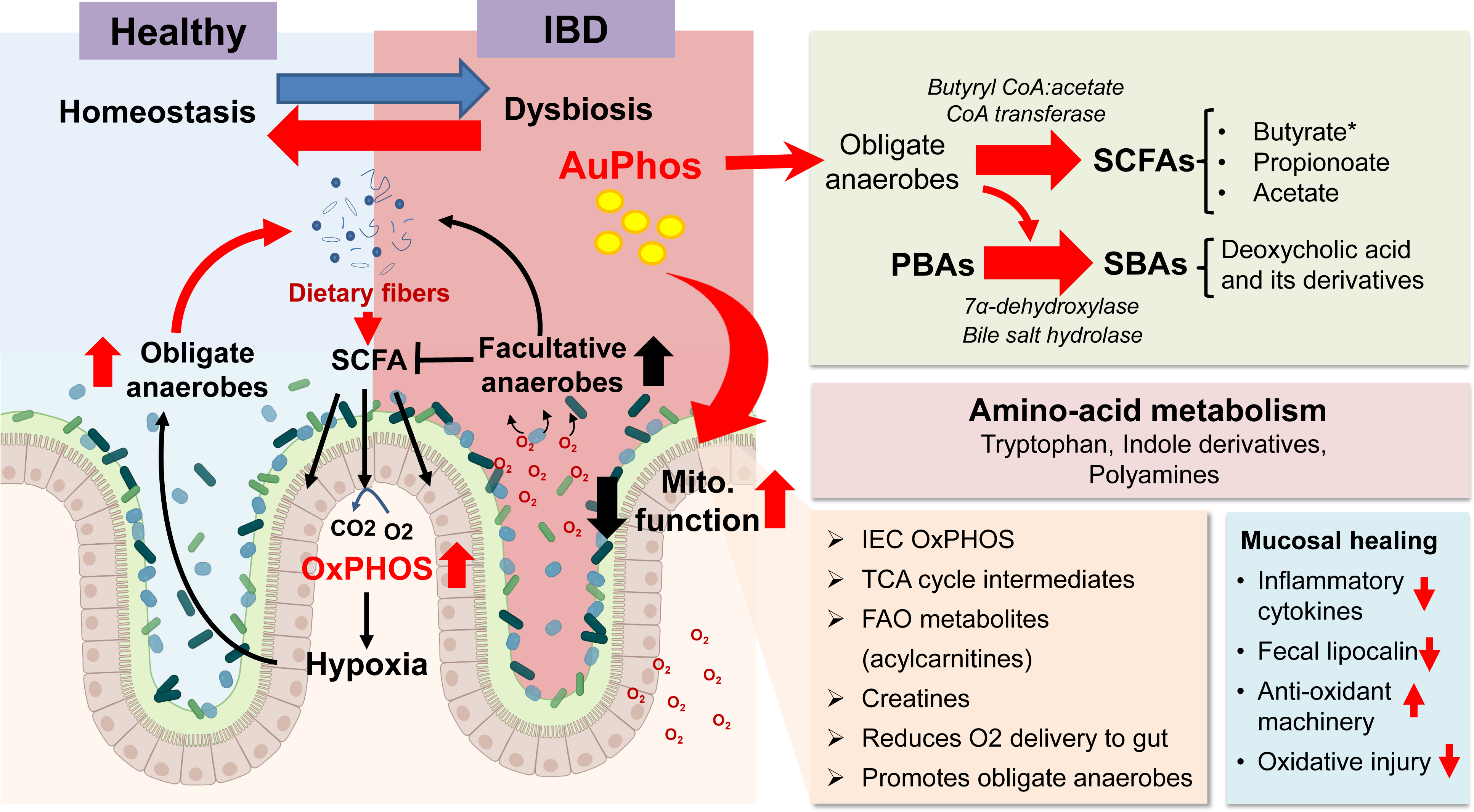

## INTRODUCTION

The intestine harbors a diverse array of microorganisms collectively termed as the gut microbiota. These microbes regulate significant metabolic, immunologic and gut protective functions. Under perturbed conditions, alterations in microbial composition, a state also referred to as microbial dysbiosis, are associated with several chronic human pathologies including inflammatory bowel diseases (IBD). Dysbiosis comprising a shift from beneficial bacteria such as *Firmicutes* and *Bifidobacteria* to an increase in pro-inflammatory bacteria such Proteobacteria, is considered a major driver of IBD pathogenesis^1–5^. Additionally, a decrease in beneficial or anti-inflammatory bacteria of certain strains such as *Bifidobacterium* and *Faecalibacterium prausnitzii* are also observed^5^. Consequentially, the microbiota diversity also decreases in IBD. A balanced gut microbiota maintains the integrity of the intestinal barrier, which prevents harmful substances and bacteria from entering the bloodstream. Dysbiosis compromises the barrier function leading to increased intestinal permeability allowing toxins and bacteria to crossover into the intestinal tissue and trigger an inflammatory response^5–7^. An aberrant immune response to certain microbes can exacerbate inflammation in the gut^7,8^. Thus, dysbiosis induces immune dysregulation that contributes to chronic inflammation in IBD. Dysbiosis-induced metabolic aberrations such as decreased short-chain fatty acids can exacerbate gut inflammation^9^. Several key studies have implicated genetic factors involved in the underlying development of inflammation^10–12^. These genes can influence a person’s susceptibility to IBD and their response to treatment. Advances in our understanding of dysbiosis has led to therapeutic innovations aimed at manipulating the gut microbiota in IBD. Probiotics, prebiotics, and fecal microbiota transplantation (FMT) are among the several strategies being explored to mitigate gut inflammation by restoring a more balanced microbiota. These interventions attempt to correct the dysbiotic microbial profile by enhancing beneficial bacteria (obligate anaerobes: e,g, *Firmicutes*) and reducing harmful pathobionts (facultative anaerobes: e,g. *Proteobacteria*) that further contribute to the inflammatory process in the colon.

Current literature supports a strong association between reduced mitochondrial function and intestinal inflammation^13–17^. Genetic studies indicate that IBD risk-conferring alleles include SNPs that alter REDOX function in patient tissue^10–12^. The causative nature of these data is supported by prospective studies in pediatric ulcerative colitis (UC) and Crohn’s disease (CD) that identify mitochondrial dysfunction as a predictor of poor clinical outcomes. There are several explanations linking mitochondrial dysfunction to IBD. Deficient mitochondrial function compromises barrier function, antioxidant defenses, stem cell gene expression, autophagy and epithelial ATP production^15,16,18–20^. More recently, data from Baumler and colleagues support a model where mitochondrial dysfunction (and reduced mucosal hypoxia) promotes disease-associated dysbiosis^2,3,21^. Their data showed that the induction of colitis coincides with a transition from a microbiota comprised of obligate anaerobes (e.g. *Firmicutes*) to one dominated by facultative anaerobes (e.g. Bacteroidetes) that utilize O2 (dysbiosis). Normally, intestines are lined with differentiated intestinal epithelial cells (IECs) with high mitochondrial function that consume O2 (O2) that is coupled to oxidative phosphorylation (OXPHOS). Epithelial O2 consumption reduces O2 availability to the microbiome and promotes an environment that supports expansion of anaerobic flora. Anaerobic flora ferment dietary fiber and produce short chain fatty acids (SCFA) that provide substrates for IEC OXPHOS^3,22–25^. Over 30 years of data support the contention that IEC in Crohn’s disease (CD) and ulcerative colitis (UC) are deficient in mitochondrial function due to oxidant injury. These data coincide with analysis of dysbiotic microbiomes in CD and UC characterized by increased blooms of facultative anaerobes that utilize O2 (e.g. *Proteobacteria*) and reduced obligate anaerobes (e.g. *Firmicutes*) that thrive in rigid anaerobic environments. Current IBD therapeutics yield suboptimal results in inducing durable remission and fail to target mitochondrial dysfunction.

In IBD, inflammation-induced damage impairs mitochondrial function that limits OXPHOS and reprograms IECs to O2 independent lactic fermentation of sugars. Reduced IEC O2 consumption increases the amount of O2 available to the microbiome that promotes expansion of facultative anaerobes that do not produce SCFA. Together these findings provide a plausible link between mitochondrial dysfunction and dysbiosis as a driver of IBD pathogenesis. Furthermore, we posit that restoration of mitochondrial metabolism contributes to IBD therapy by promoting the establishment of a microbiome dominated by obligate anaerobes.

## MATERIALS AND METHODS

### Mice

The study used 6-8-week-old C57BL/6 wildtype (WT) and *Il10-/-* C57BL/6 mice purchased from The Jackson Laboratory (Bar Harbor, MD), The mice were housed in the University of Kentucky (UKY) and Lexington VA healthcare animal care facilities under specific pathogen-free conditions. The germ-free (GF) *Il10-/-* and WT mice on 129.SvEv background were procured from National Gnotobiotic Rodent Resource Center at UNC-Chapel Hill, NC USA. The mice (7.5–10 weeks-old) were housed in a BSL2 isolator room in filtered top sterile cages. All mice were given normal chow and water ad libitum and had normal day/night cycles.

### Ethical Considerations and Human Biopsy Samples

The study was conducted at the University of Kentucky, US. Informed consent was obtained from all participants, and the study was approved by the Ethics Board of the University of Kentucky (#48678). Human intestine biopsy specimens were obtained from ulcerative Colitis (UC) and crohn’s Disease (CD) actively enrolled patients into a prospective study evaluating biopsy site healing. Biopsies from healthy control patients undergoing routine colon cancer surveillance, were obtained for comparison.

### Colitis Induction and AuPhos treatment

For the acute DSS model of colitis, 6-8-week-old WT mice were randomly selected to receive 2% dextran sodium sulfate (DSS) 40-50,000 MW (Affymetrix, Santa Clara, CA) in drinking water for 7 days followed by normal drinking water afterwards. The onset of spontaneous colitis was accelerated and synchronized by giving piroxicam-chow (Harlan Teklad, Madison, WI, standard powdered rodent chow) to 5–6-week-old C57B6 *Il10-/-* mice for 2 weeks. Piroxicam (USP grade) is procured from Sigma-Aldrich, St. Louis, MO). The mice received 65 mg of piroxicam / 250 g of chow during week 1 and 85 mg piroxicam/250 g chow during week 2. Mice were then placed on normal rodent diet without piroxicam for the remainder of the protocol. All studies and procedures were approved by the Institutional Animal Care and Use Committees of the University of Kentucky (protocol # 2019-3245) and Lexington VA healthcare research.

### In vitro growth of E. coli Nissle 1917 strains under anaerobic or microaerophilic conditions

Liquid overnight cultures of each E. coli strain were washed twice with phosphate-buffered saline (PBS), diluted to a concentration of 1 × 10^7^ CFU/ml, and mixed in a 1:1 ratio of the E. coli Nissle 1917 wild type and an isogenic mutant strain. These cultures were then diluted to a concentration of 1 × 10^6^ CFU/ml in M9 minimal medium (M9 salts medium with 0.1% glucose) and incubated statically for 24 h at 37°C in an anaerobic chamber (Shel Lab Bactron II; 5% hydrogen, 5% carbon dioxide, 90% nitrogen) or in a hypoxia chamber (Coy Laboratory Products) set at 0.8% oxygen. The competitive index (ratio of wild type/mutant) was determined by spreading serial 10-fold dilutions on LB agar plates containing the appropriate antibiotics.

### Hypoxia staining

For detection of hypoxia, 60 mg/kg of pimonidazole HCl (Hypoxyprobe-1 kit; Hypoxyprobe) was administered intraperitoneally to AuPhos- or vehicle-treated mice. After 1 h, mice were euthanized, colon extracted and swiss-roll were fixed in 10% phosphate-buffered formalin followed by paraffin block preparation. Paraffin-embedded tissues were cut into 5µ sections, which were then processed for IHC staining for pimonidazole adducts. Briefly, colon sections were blocked with mouse-on-mouse blocking reagent (Vector Labs) and probed using mouse anti-pimonidazole monoclonal IgG1 (monoclonal antibody [MAb] 4.3.11.3). Slides were then stained with FITC-conjugated goat anti-mouse secondary antibody followed by counter staining with 4’,6’-diamidino-2-phenylindole (DAPI) in Prolong Gold mountant. Representative images were obtained using a Zeiss Axioscan Z.1 fluorescence microscope.

### Fecal Microbiota Transplantation (FMT) slurry preparation

Stool samples will be collected from active endoscopically-confirmed IBD patients, under an institution-approved IRB. Freshly collected stools will be transported to the laboratory on wet ice, then aseptically aliquoted in an anaerobic chamber and stored at −80 °C^26^. Stool samples will be characterized for dysbiostic microbiome using 16S rRNA sequencing. Stool samples (100mg) with a high obligate:facultative anaerobe ratio will be mixed with sterile, preconditioned phosphate-buffered saline (1 ml) and vortexed (5min) at room temperature to prepare fecal slurries. Solids will be sedimented by centrifugation at 200 rpm for five minutes at 4 °C, and equal volume of supernatants from individual samples will be combined to make IBD stool cocktail. Aliquot (500 µL) of this cocktail will be prepared in anaerobic conditions and stored in −80 °C until colonization.

### Colonization of GF mice gut with FMT cocktail

Frozen FMT slurry (500 µL; 100mg/mL) prepared from human IBD donors were thawed in anaerobic chamber and diluted to 10mg/mL with pre-reduced sterile PBS (addition of 4500uL of sterile PBS to 500uL of 100mg/mL slurry to generate 5000uL of 10mg/mL slurry). Each GF-WT (129-strain WT) mouse was gavaged with 2mg of FMT cocktail, which corresponds to 200uL of 10mg/mL slurry.

### Fecal Lipocalin 2 Assay (f-LCN2)

Fresh fecal samples were collected biweekly from the experimental mice and frozen immediately in dry ice to retain the microbial integrity and stored at −80 °C. Fecal lipocalin-2 (f-LCN2) levels were measured by R&D Systems’ ELISA kit, according to the manufacturer’s protocols99. To quantify the f-LCN2 values, 10–20 mg frozen fecal samples were dispersed in PBS containing 0.1% Tween 20 and incubated overnight at 4 °C. The slurry were then briefly vortexed to obtain a homogenous fecal suspension and centrifuged for 10 min at 12,000 rpm and 4 °C. f-LCN2 concentrations were determined in the collected supernatants using the color reagents of hydrogen peroxide and tetramethylbenzidine at 450 and 570 nm by a microplate reader (Biotek Synergy HT, BioTek®Instruments, Inc.,Winooski, VT, USA).

### Histologic Assessment of Colitis

Entire colons were opened longitudinally and rolled up onto a syringe plunger. The tissue was fixed in 10% neutral buffered formalin for 24-48h, removed from plungers without unrolling, and processed for paraffin-embedding followed by subsequent sectioning. Then 5 µm longitudinal sections were cut and stained with hematoxylin and eosin (H&E) for histologic assessment. Inflammation was scored on a scale from 0–4 per the scoring system described by Berg et al. Briefly, the mucosal inflammation in the proximal colon, distal colon, and rectum were blindly assessed according to a scoring system ranging from 0–4 for the degree of lamina propria and submucosal mononuclear cellular infiltration, crypt hyperplasia, goblet cell depletion, and architectural distortion. Histological scores are represented individually and as a sum of scores (maximum score = 12) for the entire large intestine, as previously validated.

### Gut microbiota 16S rRNA sequencing and profiling

Total bacterial DNA was extracted from frozen (−80°C) fecal samples from AuPhos- or vehicle treated mice using PowerFecal DNA/RNA Kit (Qiagen, Hilden, NRW, Germany), according to the manufacturer’s instructions. The purity and concentration of bacterial DNA were determined using the NanoDrop system (Thermo Fisher Scientific, Madison, WI, USA). The samples were then prepared for 16S rRNA sequencing. The V4–V5 regions of 16S rRNA gene were amplified using specific primers and libraries were constructed using the Hyper Library Preparation Kit from Kapa Biosystems (Roche Diagnostics, Indianapolis, Indiana, USA), followed by Qubit verification. Libraries were then pooled at equimolar concentrations and high-throughput sequencing was performed by Illumina HiSeq 6000 high-throughput sequencer (Illumina, San Diego, USA). Differential abundance analyses were performed using ANCOM (Analysis of composition of microbiomes) with Holm–Bonferroni multiple-testing correction. The Shannon index was calculated to measure alpha diversity (within-group diversity), and the pairwise difference in alpha diversity tested using a one-way ANOVA test.

### Real-time Quantitative Reverse-Transcription PCR

Total RNA was isolated from Allprotect segments of the intestine or sorted cells using the RNeasy Mini Kit (Qiagen, Valencia, CA) and reverse transcribed using the High Capacity cDNA Reverse Transcription Kit (Applied Biosystems, Foster City, CA). Individual genes were run on an ABI Step One Plus Real Time system using Power SYBR green PCR master mix (Applied Biosystems). Primers were designed by Primer Express software 3.0 (Applied Biosystems) based on nucleotide sequences from the National Center for Biotechnology Information data bank. Multiple genes were run for normalization of data (GAPDH, β-actin, β2M and Hsp90, and/or the 18S and 28S rRNAs) and used as an internal reference. All assays were performed in triplicate and fold changes were calculated using the ΔΔCT method.

### Histology and immunohistochemistry (IHC)

Tissues were fixed in 4% neutral buffered formalin overnight, processed through paraffin and sectioned at 5 μm. For laboratory staining of vimentin, paraffin sections were rehydrated through graded alcohols and antigen retrieval was performed using Target Retrieval Solution (Dako, Carpinteria, CA, United States), pH 6.0, in a decloaking chamber. Sections were incubated with NDUFB6, MTCO1 and COX5A antibodies (Invitrogen) followed by anti-rabbit or anti-mouse peroxidase-labeled polymer (Dako). Sections were developed using 3,3’-diaminobenzidine (DAB) tetrahydrochloride chromagen (Dako) and counterstained with hematoxylin.

## RESULTS

### AuPhos-induced mitochondrial function promotes healthy anaerobic microbiome in mice

This study investigated the ability of a novel AuPhos compound^27^ to create luminal micro-anaerobic environment inside the gut, which promotes the expansion of healthy obligate anaerobic bacteria. Studies here test the notion that AuPhos-improved mucosal mitochondrial function enhances mucosal O2 utilization thereby reducing O2 delivery to the enteric microbiome. Reduced enteric O2 availability shifts microbial metagenomics, meta-transcriptomics and metabolism resulting in increased production of short-chain fatty acids (SCFA) amongst other predicted changes [e.g. bile salt, amino-acid (tryptophan, alanine), FAO, TCA cycle metabolites]. We have earlier shown that AuPhos has a potential to increase the mitochondrial function in both in vitro and in vivo systems^28–30^. To understand whether the changes in gut microbial communities are due to the direct effect of AuPhos on mucosal metabolism rather than effect of resolution of inflammation, we treated uninflamed WT mice with AuPhos. Healthy uninflamed WT mice showed differential beta-diversity of bacteria in stool samples from AuPhos-fed (2.5- and 250mg/Kg; q3d; 2 weeks) mice compared to vehicle control mice, as revealed on Principal Coordinates Analysis (PCoA) plot (**Fig. 1A**). Metagenomic (16S rRNA) analysis showed reduction in relative abundance of (O2 consuming) *Proteobacteria*, which includes facultatively anaerobic *Enterobacteriaceae* family, in AuPhos-fed mice (**Fig. 1B-C**). Conversely, AuPhos treatment dose-dependently increased the relative abundance of signature obligate anaerobic bacteria, *Firmicutes* (**Fig. 1B).** At the genus level, AuPhos treatment increased the relative abundance of *Clostridium*, *Biophila*, *Odoribacter*, *Parabacteroides* and *Bacteroides* in a dose-dependent manner (**Fig. 1C**). PICRUST2 and LDA revealed that AuPhos decreased bacterial LPS biosynthetic pathway and increased overall fatty acid biosynthesis pathways (**Supplementary Fig 1**). AuPhos increased fecal levels of SCFAs with >35% increase in butyrate and propionate levels, and 60% increase in compared to stools from vehicle-treated mice (**Fig.1D-E**). Next, we tested the AuPhos potential to modulate microbial shift under inflamed conditions using DSS-induced colitis. Treatment of DSS-induced colitis mice with AuPhos (25mg/kg, q3d) decreased the relative abundances of bacteria that belongs to *Proteobacteria* phylum with concomitant increase in *Firmicute* phylum at the end of 3 weeks (D21) compared to vehicle control (**Fig. 1F**). Similar to uninflamed healthy mice, DSS colitis mice also showed an increase relative abundance of genus; *Clostridium*, *Biophila*, and *Odoribacter*, after D21 treatment with AuPhos compared to vehicle control (**Fig. 1G**). Earlier we have shown the potential of AuPhos in mitigating inflammation, improving colitis score and barrier function in DSS-colitis mice^28^. These results indicate the potential of AuPhos in expanding the microbial population of obligate anaerobes that would ferment dietary fibers into SCFAs, which provide energy substrate to intestinal epithelial cells (IECs) and further advances ulcer healing process under colitis conditions.

**Figure 1.**
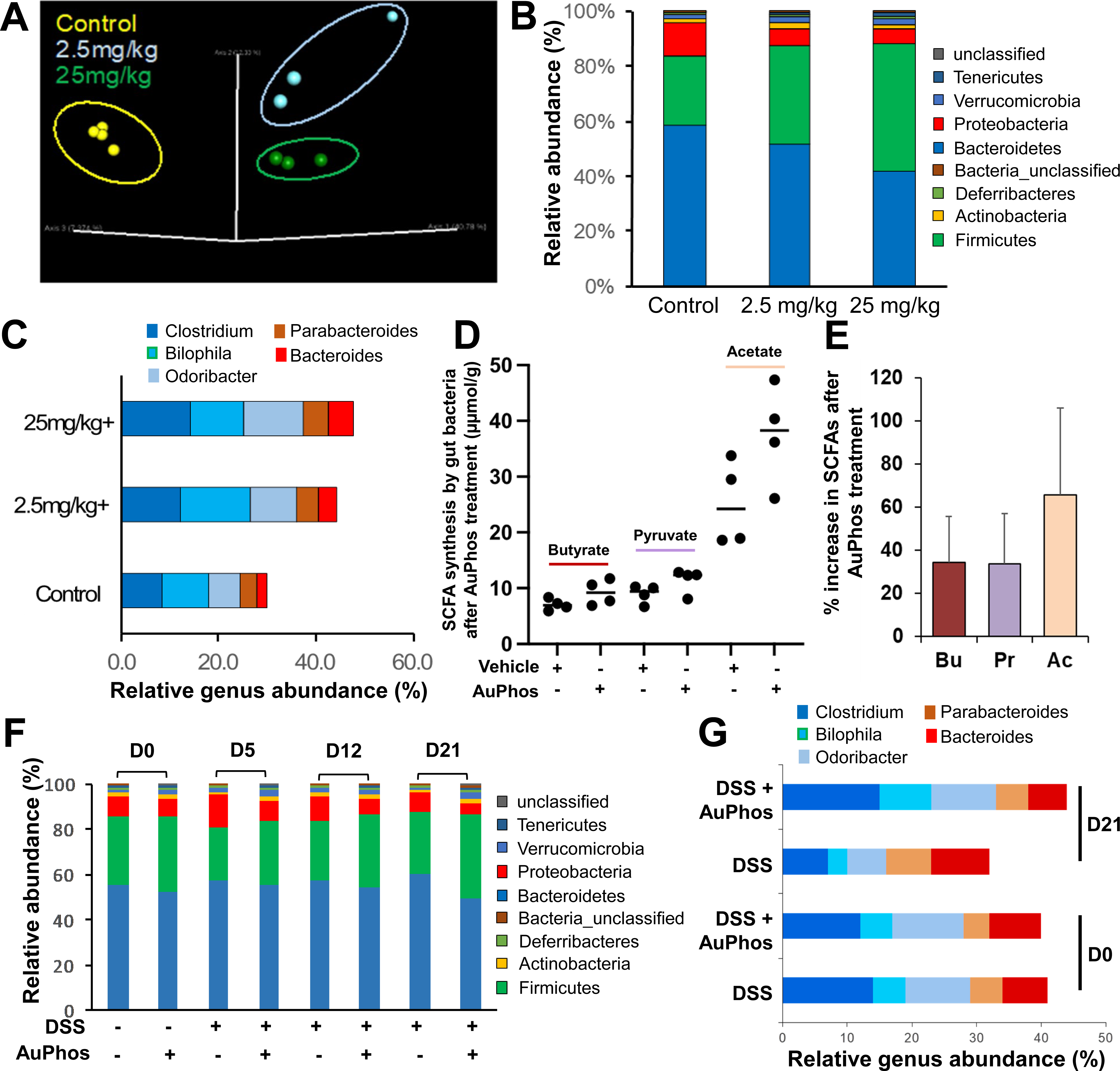
Orally administered AuPhos promotes the growth of obligate anaerobes in B6 WT mice. A. Principal Coordinates Analysis (PCA) plot showing beta-diversity based upon the unweighted UniFrac distances, where each point represents a single sample. Healthy uninflamed WT mice were orally treated (q3d, 2 wk) with vehicle (control, Yellow), AuPhos 2.5 mg/kg (Blue) and AuPhos 25 mg/kg (Green). Relative abundance of **B.** gut bacterial taxa at phylum level, and **C.** predominant bacterial genus (>5%) in the stool samples of treated WT mice. **D.** Effect of AuPhos on SCFA levels in the stool samples from vehicle control and AuPhos-fed (25mg/kg) WT mice along with percentage increase in SCFAs in AuPhos vs vehicle in **panel-E**. Short chain fatty acids were determined by ^1^H NMR and normalized to frozen fecal pellet weight. Bu=butanoate Pr=propanoate, Ac =acetate. AP = AuPhos treated mice. **E.** Relative abundance of gut bacterial taxa at phylum level and **F.** predominant bacterial genus (>5%) in the stool from DSS-mice (+/- AuPhos). Mice were allowed free access to food and drinking water containing 2% (wt/vol) DSS for 7 days, and recovery was allowed for an additional 2 weeks.

### AuPhos increases O2 utilization by surface epithelial cells and creates hypoxic microenvironment

In order to test the notion that increased abundance of obligate anaerobes in AuPhos treated mice is due to luminal O2 deprivation, we performed experiments to assess the bioavailability of enteric O2 in the colon of WT and DSS-colitis mice. Direct O2 measurements in the gut lumen is difficult because facultative anaerobic bacteria readily consume O2. We thus assessed the bioavailability of oxygen using indicator strains of *E.coli* that were either aerobic respiration proficient (*E. coli* Nissle 1917 wild type) or aerobic respiration-deficient under microaerophilic conditions [*E. coli* Nissle 1917 with cytochrome oxidase (*cydA)* mutant]. AuPhos-induced anaerobiosis was tested by measuring the effect of AuPhos in modulating the growth of WT and cydA mutant strains. Orally administered AuPhos promotes the growth of cydA mutant *E.coli* that display respiratory properties akin to obligate anaerobes, in both healthy uninflamed WT (**Fig. 2A**) and DSS-colitis (**Fig. 2D**) mice, inoculated with a 1:1 mixture of both bioindicator strains. AuPhos treatment significantly increased the number of cydA mutant and concertedly deceased WT *E.coli* recovered from colonic contents of WT (**Fig. 2A**) and DSS-colitis (**Fig. 2D**), compared to vehicle treated groups. The fitness advantage conferred by aerobic respiration was assessed between vehicle- and AuPhos-treated mice by determining the ratio of WT vs. cydA mutant *E.coli* (competitive index; CI) recovered from colonic contents. AuPhos significantly reduced the CI in WT (**Fig. 2B**) and DSS-colitis mice after day-7 of treatment (**Fig. 2E**), which supported our notion that AuPhos decreased O2 bioavailability in the intestinal lumen. Evidences from others^3^ and our earlier work suggest that colonic anaerobiosis is established and maintained by actively respiring (high OXPHOS) colonic surface epithelial cells. High O2 consumption capacity of luminal colonocytes maintain a state of physiological hypoxia at the epithelial surface.

**Figure 2.**
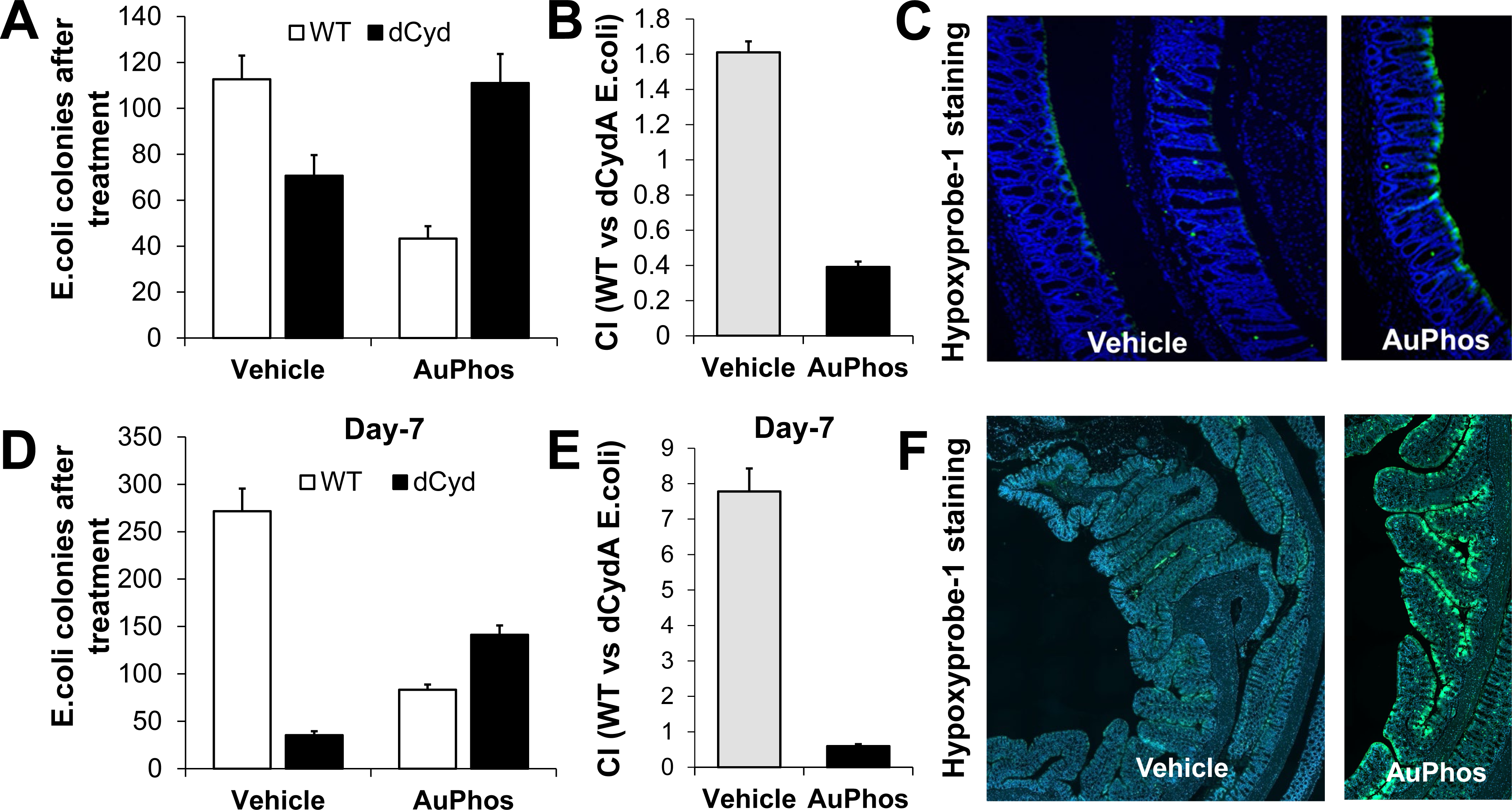
Orally administered AuPhos increases oxygen utilization by surface IECss and decreases epithelial oxygenation. Effect of AuPhos on enteric oxygenation in uninflamed WT (**A-C**) and DSS colitis (**D-F**) mice. Numbers of WT and cydA mutant strains of *E. coli* bacteria recovered from colonic contents of vehicle or AuPhos-fed WT (**panel-A**) and DSS mice at D7 of post-treatment (**panel-D**). The fitness advantage conferred by aerobic respiration was assessed by determining the competitive index (CI) of *E. coli* wild type (wt) and a cydA mutant recovered from colonic contents of WT (**panel-B**) and DSS mice (**panel-E**). AuPhos-induced hypoxia in surface IECs was assessed by hypoxyprobe staining. Binding of pimonidazole was detected using Hypoxyprobe-1 primary antibody and FITC-conjugated goat anti-mouse secondary antibody (green fluorescence) in colon sections counterstained with DAPI nuclear stain (blue fluorescence) in AuPhos-fed WT (**panel-C**) and DSS (**panel-F**) mice compared to vehicle control.

AuPhos-induced hypoxia in surface IECs was assessed by Hypoxyprobe-1 staining in sections from pimonidazole HCl-infused mice, which is reduced to hydroxylamine intermediates under hypoxic conditions and form adducts with DNA or protein (**Fig. 2C,F**). Pimonidazole staining revealed that the colonic surface of AuPhos-fed WT (**Fig. 2C**) and DSS-colitis (**Fig. 2F**) mice had comparatively higher hypoxia than vehicle-treated mice, suggesting that AuPhos decreased epithelial oxygenation.

### AuPhos induced ulcer healing in chronic colitis in *Il10-/-* mice

Acute colitis mouse models of IBD fail to mirror the chronic nature of the disease, because these are self-limiting models and inflammation spontaneously resolves by itself with few days. We, therefore, utilized piroxicam (Px)-accelerated *Il10-/-* mice for rapid, uniform and sustained development of colitis^31,32^, which has earlier been used by our research group^32^. Looking at the positive effect of AuPhos in regulating microbiome and correcting dysbiosis, we wanted to assess the AuPhos potential to heal ulcers in chronic Px-accelerated *Il10-/-*. Piroxicam administration for 2 weeks accelerates and synchronizes the onset of colitis in *Il-10−/−* mice with marked colonic crypt hyperplasia, muscle wall thickening, mononuclear cell infiltration of the LP, and transmural inflammation. Nearly all Px-*Il-10−/−* mice (>90%) developed epithelial ulceration and significantly higher inflammatory scores after 2 weeks compared to piroxicam-treated B6 controls^32,33^. Treatment with AuPhos (25mg/kg; q3d), initiated at D28 (based of fecal LCN2 levels; data not shown), significantly decreased the disease activity (p<0.05; **Fig. 3A**) and fecal inflammatory marker LCN2 (p<0.05; **Fig. 3B**) at D55 (end of treatment) compared to vehicle-treated group. AuPhos treatment of Px-*Il10-/-* mice repaired mucosal ulceration and showed no significant inflammatory infiltrate or hyperplasia (**Fig. 3C**). AuPhos significantly decreased overall colitis score and severity score in Px-*Il10-/-* mice (**Fig. 3D**) despite the aggressive nature of the transmural colitis. Further assessment of pro-inflammatory markers, showed significant decrease in mRNA levels of *Il-1β*, *Il-6*, *IFN-γ*, *Il-17*, *Il-18*, and *IP-10* (**Fig. 3E**) compared to vehicle group. In order to assess whether the AuPhos-resolved inflammation parameters in Px-*Il10-/-* mice are correlated with mitochondrial function, we measured the transcripts level of mitochondrial complexes. AuPhos significantly increased the mRNA levels of mitochondrial complex I (*ND5*) and complex V (*ATP6*) at end of the treatment compared to vehicle group (**Fig. 3E**). Taken together, these results suggested that AuPhos has a potential to attenuate severe colitis in *Il10-/-* mice.

**Figure 3.**
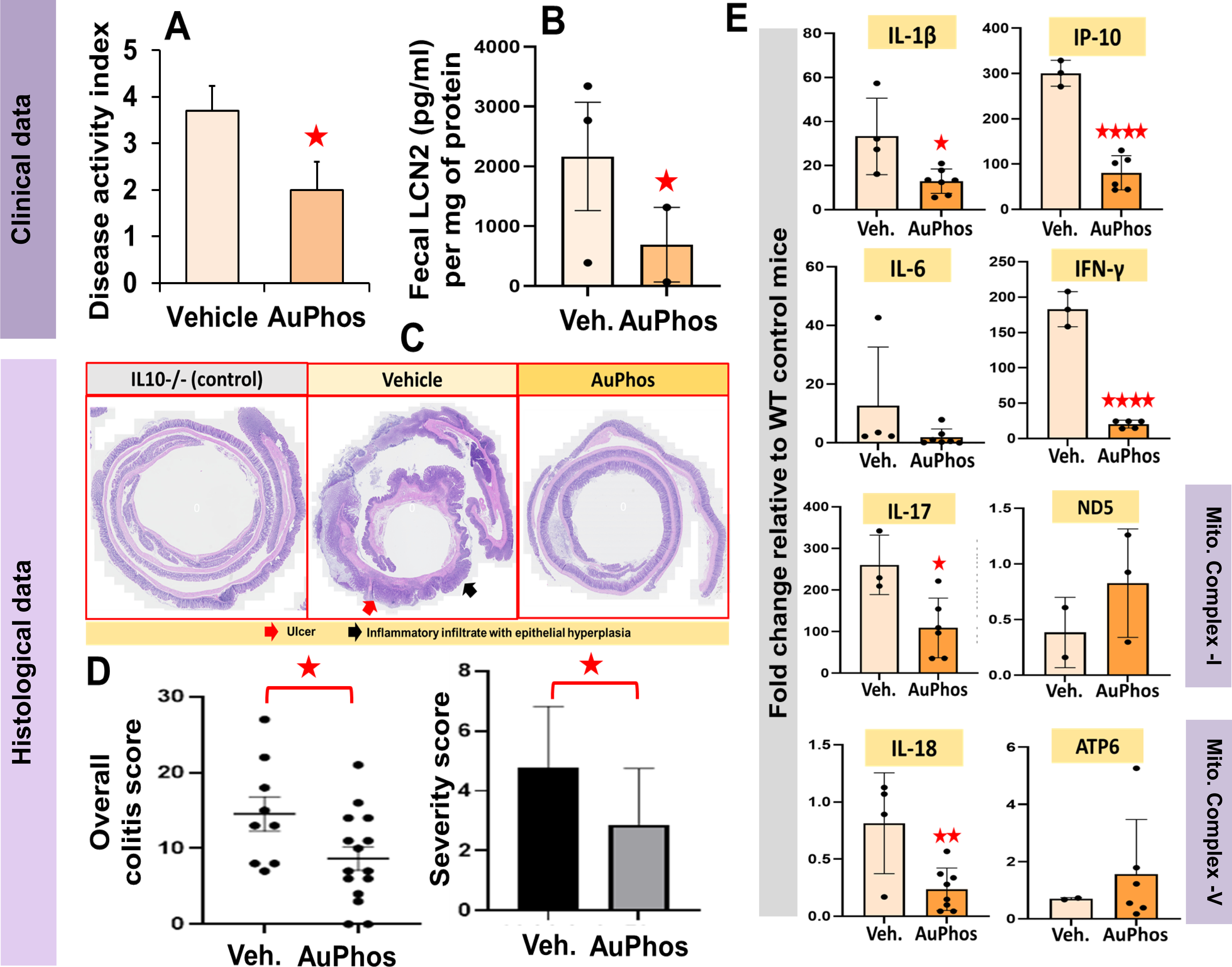
AuPhos-induced ulcer healing in chronic colitis in IL10-/- mice. AuPhos treatment (D28-D55) significantly (*p<0.05) decreased (A) disease activity index and (B) fecal lipocalin-2 (LCN2) in the stool samples collected at D-55 from Px-IL10-/- vs.vehicle-treated mice. C) Decreased severity of colonic inflammation in AuPhos-treated Px IL10-/- mice at D56. Red arrow represents areas of colonic ulceration and black arrow represents inflammatory infiltrate with epithelial hyperplasia. D) AuPhos significantly decreased overall colitis score and severity score in Px IL10 KO mice. E) RT-qPCR shows significant reductions in mRNA for inflammatory cytokines and chemokines with concomitant increase in transcripts of components of mitochondrial complexes at D56 in AuPhos-fed mice. Statistical significance was determined by unpaired t-test. *p < 0.05; **p < 0.01; ***p < 0.001; ****p < 0.0001.

### AuPhos induces expression of mitochondrial complexes, stem cell markers and reduces inflammatory markers in human colonic biopsies

AuPhos effect on reducing inflammation parameters and upregulating mitochondrial complexes in Px- *Il10-/-* mice, encourage us to investigate its direct role on colonic mucosa in the absence of microbiome. Transcriptomic analysis was performed to assess the role of AuPhos in modulating markers of mitochondrial complexes, stem cells and inflammation in ex vivo treated human colonic biopsies from normal control, UC, and CD patients. AuPhos (0.5 uM) treatment (3h) of human colonic biopsies from normal control patients significantly increased the mRNA level of *Ndufa4* (∼2-fold; p<0.05), *Ndufb6* (∼2-fold; p<0.05), and *Cox5a* [∼1.5 fold; not significant (ns)] compared to vehicle treatment (**Fig. 4A**), indicating the AuPhos effect on enhancing mitochondrial complexes under uninflamed conditions. Interestingly, ex vivo treatment of biopsies from IBD patients also showed significant upregulation of transcripts of *Ndufa1* (UC: ∼4-fold; p<0.05, CD: ∼3.7-fold; p<0.05), *Ndufb4* (∼1.8-fold; p<0.05, CD: ∼2.5-fold; p<0.01), *Ndufb6* (∼1.66-fold; p<0.05, CD: ∼2.7-fold; p-value: ns) and *Cox5a* (∼5.1-fold; p<0.01, CD: ∼1.6-fold; p-value: ns), compared to median fold-change values of respective vehicle controls (**Fig. 4A**). Further, IHC analysis of NDUFB6 staining also showed significant increase in mitochondrial complex I protein after treatment (3h, ex vivo) of human colonic biopsies of normal (∼1.64-fold; p<0.05) and UC (∼2.72-fold; p<0.05), compared to vehicle group (**Fig. 4B,C**). Interestingly, AuPhos also increased mRNA for intestinal stem cell (ISC) markers (Lgr5) and reduced mRNA for inflammatory cytokine (IL1β, MCP-1, RankL, TNF) in biopsies from mild UC patients (**Fig 4D,E**). Effects on ISC markers are consistent with prior data showing that OxPhos promotes Wnt signaling in small bowel ISC (ref 47 grant).

**Figure 4.**
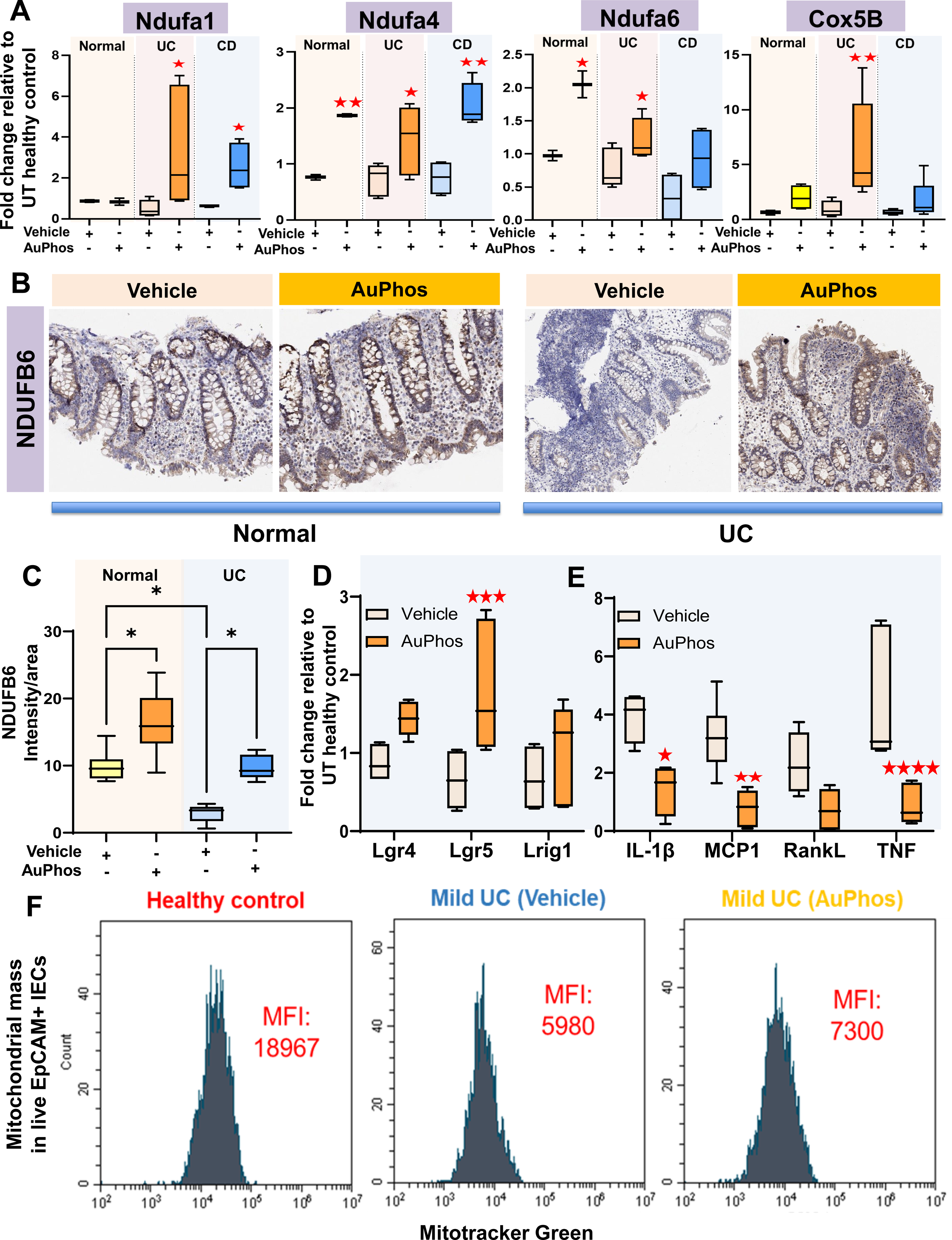
AuPhos treatment (0.5uM) of human IBD colonic biopsies significantly increases. **A.** mitochondrial complex I (**Ndufa1, Ndufa4, Ndufb6**), complex IV (**Cox5B**) transcripts, **B.** IHC staining of Ndufb6 in normal and UC, along with imageJ analysis showing **C.** intensity per unit area **D.** stem cell (**Lgr5**) gene mRNA and **E.** decreases cytokines/chemokines mRNA (**IL-1β, MCP1, RankL, TNF**) compared to vehicle controls. **F.** Flow cytometric assessment of mito mass (mitotracker green) in live EpCAM+-sorted IECs from healthy controls and mild UC colonic biopsies. In summary, AuPhos-treatment (0.5uM) increases mitochondrial mass in IECs from mild UC patients after 3h treatment compared to vehicle control. Statistical significance was determined by 2-way ANOVA using Sidak’s multiple comparison; *p < 0.05; **p < 0.01; ****p < 0.0001.

Flow cytometric assessments of mitochondrial mass in live (EpCAM+-sorted) IECs from healthy controls and mild UC biopsies showed that **AuPhos** increases mitochondrial mass in normal and UC (**Fig 4F**). Data from human IBD patients also suggest the potential of AuPhos as a novel mitochondria-targeted oral therapeutic for IBD patients.

### AuPhos corrects dysbiotic changes in germ-free *Il10-/-* mice reconstituted with human IBD stool

Establishment of sustained inflammation in Px-*Il10-/-* mice provided us a platform to investigate the efficacy of AuPhos in ameliorating the chronicity of inflammation akin to those present in severe UC patients. However, the gut microbiome of these *Il10-/-* mice does not reflect the true dysbiotic enteric environment as present in the IBD patients. We, therefore, performed investigations to identify the ideal human fecal microbiota transplant (FMT) mouse model that could mimic the dysbiotic microbial imbalance in human IBD gut. Briefly, conventional microbiota from healthy donor were transplanted into germ-free (GF) B6 WT (n=6) and *Il10-/-* mice (n=6) (**Fig. 5A**) and 16S rRNA sequencing was performed on the stool samples after two weeks post-colonization. Our analysis of fecal 16s rRNA-based microbial profiling showed that relative abundance of *Enterobacteriaceae* (mainly *Protobacteria*) increases in *Il10^-/^*^-^ mice reconstituted with conventional gut microbiota compared to WT B6 (**Fig. 5A**). Similar increase in *Protobacteria* was observed when human IBD stool from three IBD patients were transplanted into GF-*Il10^-/^*^-^ mice (129.SvEv background) compared to 129.SvEv WT mice (**Fig. 5B**). Data from figure 5A and B suggests that *Il10-/-* mouse strains are excellent models to study gut dysbiosis mimicking microbial imbalance in human IBD gut. Given the likelihood that **AuPhos** increases mitochondrial functions of IEC and reverses DSS-induced inflammation, we tested whether **AuPhos** modulates gut microbiota in both healthy (uninflamed) and humanized germ free (**HuGF**) *Il10*^-/-^ colitis mice reconstituted with Hu IBD stool (**Fig. 5C-H**). Untreated **HuGF-*Il10*^-/-^** mice showed significant increases in relative abundance of *Proteobacteria* after two weeks post-colonization with human IBD stool, compared to background WT (129.SvEv) mice (**Fig. 5B**). To examine the impact of enhanced mucosal mitochondrial function (**AuPhos**) on gut microbiota in colitis, we took advantage of the **HuGF** 129 IL-*Il10^-/-^* and **HuGF WT** mice housed at the UNC’s National Gnotobiotic Rodent Resource (NGRC) facility (we procured these **GF** mice and reconstituted with **Hu** IBD stool in our VA facility). We utilized **HuGF-*Il10*^-/-^** mice to develop spontaneous colitis and treated with **C** or **AuPhos** started after **HuGF-*Il10*^-/-^** mice developed severe colitis at D28, based on f-LCN2 assay (**Fig 5D**). The mice were treated with **AuPhos** (i.g.; 10mg/kg, q3d) for over 2 weeks and assessed on D46. **AuPhos** significantly reduced the level of fLCN2 (**Fig 5D**). Histological investigation also showed significant reduction in colitis score (**Fig 5E,F**). Immunohistochemical (IHC) staining of mitochondrial complex I (NDUFB6) and complex V (MTCO1 and COX5A) on colonic sections from these **HuGF-*Il10*^-/-^** mice showed significant increase in these complexes in **AuPhos**-treated mice (**Fig 5C**). 16S rRNAseq analysis revealed **AuPhos** significantly reduced relative abundances of (O2 consuming) facultative anaerobic *Proteobacteria* and increased blooms of *Firmicutes* (obligate anaerobes) (**Fig 5G**). Importantly, we found that **AuPhos** also corrected dysbiotic microbiomes in WT mice reconstituted with Hu IBD stool (**Fig 5H**). These latter results show the effects of AuPhos in the absence of colitis.

**Figure 5.**
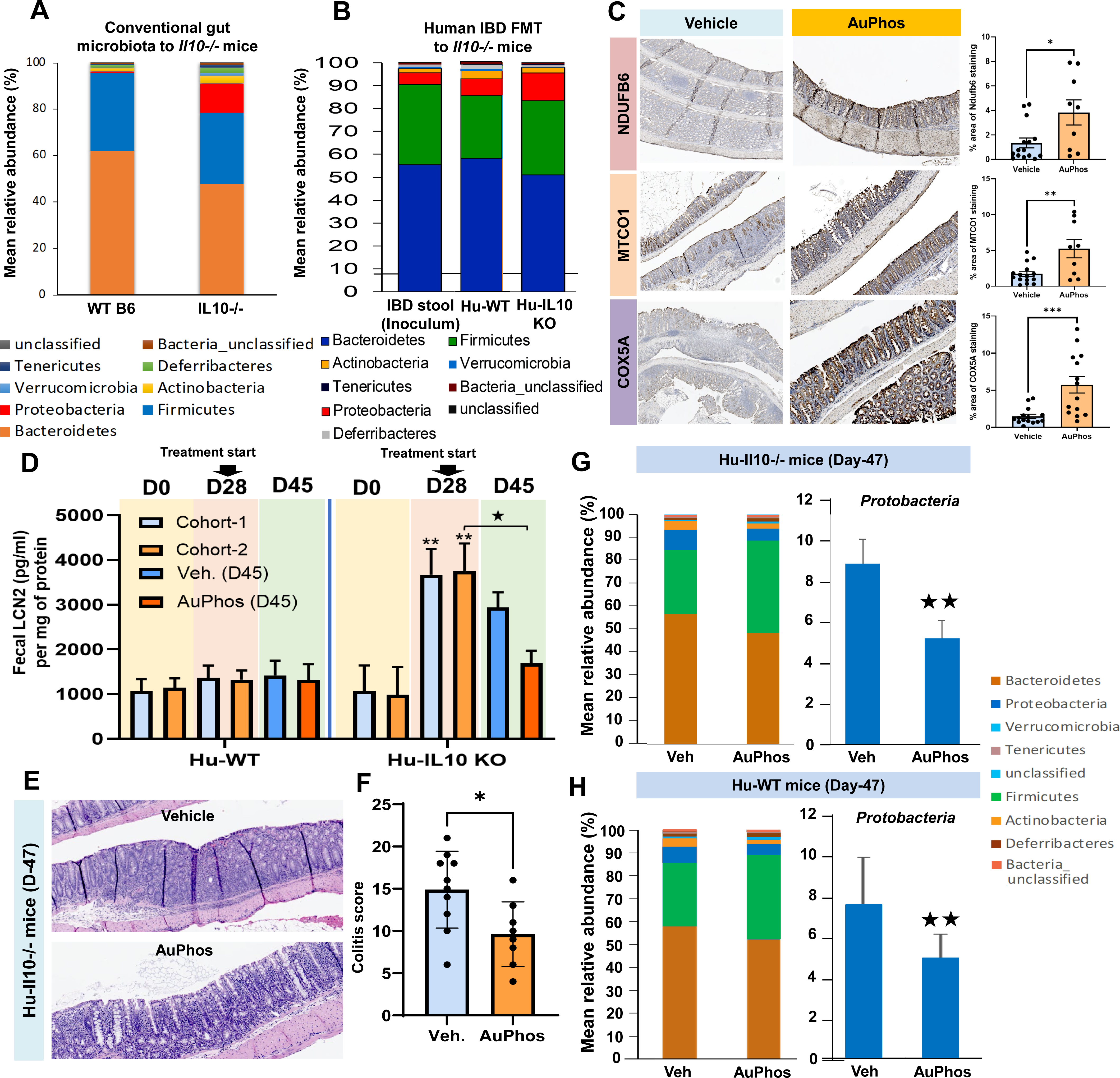
Enhanced mitochondrial function corrects dysbiosis in Hu*-Il10-/*-mice. Data in **Panel-A and –B** suggesting that *Il10-/-* mouse strains are an excellent models to study gut dysbiosis mimicking microbial imbalance in human IBD gut. Relative abundance of *Enterobacteriaceae* (mainly *Protobacteria*) increases in **A.** *Il10-/*-mice reconstituted with conventional gut microbiota compared to WT B6, and **B.** GF*-Il10-/*-mice reconstituted human IBD stool (Hu*-Il10-/*-mice) compared to Hu-WT. Graph shows alterations in the relative abundances of bacterial phyla two weeks post-colonization (n=6 mice).**C.** Comparative immunohistochemical analysis for complex I (NDUFB6) and complex IV (MT-CO1 and COX5A) staining in colon sections isolated from AuPhos- or vehicle (Veh.) treated Hu-IL10-/-. IHC staining for NDUFB6, MTCO1 and COX5A shows significantly increase in expression of these mitochondrial complexes in the AuPhos-treated mice compared to vehicle treated groups. Image processing package of ImageJ (Fiji) was used to perform DAB-intensity analysis utilizing ten non-overlapping region-of-interest (ROI) from each mouse. **D.** Fecal lipocalin-2 (f-LCN2) assessment (D0 onwards) in the stool samples from Hu-IL10-/- and Hu-WT mice. AuPhos treatment was initiated at D28 when disease is severe in Hu-IL10-/- based on f-LCN2 levels. **E.** Tissue histology and **F.** colitis score in Hu-IL10-/- mice at D47 showing significant reduction in inflammation in AuPhos-fed mice compared to vehicle group. **G & H.** Effect of AuPhos on human gut microbiota composition in **Hu-*Il10-/-*** and **Hu WT** mice. Taxonomic profiles at the phylum level for AuPhos (**G**) or vehicle-fed (**H**) mice at D47 (q3d; 2 weeks) are shown as mean relative abundance determined by HTS of 16s rRNA. **Right panel:** Mean relative abundance of proteobacterial phyla. Statistical significance was determined using unpaired t-test. *p < 0.05; **p < 0.01; ***p < 0.001

### AuPhos-enhanced epithelial mitochondrial function corrects dysbiotic metabolomics in Hu-*IL10-/-* mice

Oxygen consumption reduces O2 availability to the microbiome and promotes a healthy anaerobic environment^34^. During IBD, colitis-induced oxidant injury impairs mitochondrial function thereby limiting OxPhos and converting IECs to a glycolytic metabolism that consumes less O2^3,34^. Dysbiotic metabolites negatively affect host metabolism and immunity. Given the potential that our novel compound AuPhos upregulates IEC mitochondrial function^28,35^, attenuates colitis and corrects dysbiosis in HuGF-Il10-/- mice as shown in figure 5, we investigated the effect of AuPhos in rewiring the IBD-associated dysbiotic metabolism using untargeted metabolomics on stool samples collected from AuPhos- or Control (C)-treated HuGF-Il10-/- mice and HuGF-WT mice (**Fig 6,7**). AuPhos effects on microbial metabolites were determined in stool samples from HuGF WT and HuGF-Il10-/- mice. Mice were treated orally with AuPhos (10-mg/kg; q2d; 2wk) or vehicle (water), stool samples collected thrice a week and frozen for f-LCN2 assay (Fig 5D,F), 16S rRNAseq (Fig 5G,H) and metabolomic studies (Fig 6,7). Frozen stools were analyzed for global metabolomics and samples prior to extraction. Peak intensities were analyzed after sample-wise sum normalization followed by log (base 10) transformation and pareto scaling (R package MetaboAnalystR). The mass and retention time data were searched against the internal metabolite library, and known metabolites mapped to KEGG IDs. AuPhos significantly altered metabolites belonging to organic acid, lipid, amino acid (AA), polyamine and carbohydrate class. AuPhos altered the metabolites associated with butyrate synthesis in HuGF-Il10-/- mice (**Fig. 6A**). AuPhos significantly increased pyruvate (log2FC:2.7), succinate (log2FC:0.35), 4-hydroxybutyrate (log2FC:0.4), 2-hydroxyglutarate (log2FC:2.5), and butanoic acid (butyrate) compared to vehicle treated mice, suggesting an upregulation of pyruvate, 4-aminobutyrate (4Ab) and glutarate pathways of butyrate production (**Fig. 6A**). AuPhos also marginally increased the level of propionate and its derivates (**Fig. 6A**). Another important feature of commensal gut bacterium is the conversion of primary BAs (PBAs) to secondary BAs (SBAs), which is mediated by the enzymes associated with dihydroxylation and deconjugation^36^. AuPhos reduced IBD-associated PBAs (Ketochenodeoxycholic, Glycochenodeoxycholic Ursocholic acid, Cholate, Muricholic acid, and different derivatives of these PBAs) with concomitant increase in SBA (Tauroursodeoxycholic acid, TUDCA) (**Fig. 6B**). Intestinal inflammation also significantly alters amino acid metabolism that correlates with disease severity^37^. Our untargeted metabolomic analysis showed that AuPhos increased levels of fecal TRP (N-BOC-L-Tryptophan: 0.575, N-BOC-L-Tryptophan: 0.677, N-BOC-L-Tryptophan: 0.728, N-BOC-L-Tryptophan: 0.712, L-Tryptophan: 0.572, α,β-Didehydrotryptophan) and various indole-derivatives (3-methyldioxyindole, Indole, 3-methyloxindole, 5-hydroxyindoleacetate, indole-3-carbonyl nitrile) in AuPhos-treated compared to C-treated HuGF-Il10-/-mice (**Fig. 6C**). AuPhos treatment also reduced the level of KYN (L-Kynurenine) and its hydroxy-derivatives (3-Hydroxykynurenine) in AuPhos-treated compared to C-treated HuGF-Il10-/-mice (**Fig 6C**). There is a strong connection between increased KYN levels and IBD-associated depression 59. Besides tryptophan metabolism, AuPhos treatment significantly increased the fecal levels of arginine (L-Arginine: 0.909, L-Arginine+Cl: 0.883, L-Arginine: 0.871), ornithine (L-Ornithine, L-Ornithine-H2O: 0.737, L-Ornithine-NH3: 0.991, Ornithine: 0.904), N-Carbamoylputrescine and putrescine (N-Acetylputrescine: 0.941, Putrescine: 0.833, Putrescine-NH3: 0.857) compared to vehicle group in HuGF-Il10-/- mice (**Fig. 6D**). Our metabolomic data also showed increased fecal level of double acetylated form spermine (N1,N12-Diacetylspermine), which is a catabolic product of spermine for inter-conversion back to spermidine or putrescine^38–40^. Besides putrescine and spermine, **AuPhos** did not show much effect on level of spermidine and cadaverine in HuGF-*Il10*^-/-^ mice **(Fig. 6D).** Our untargeted metabolomics also showed increased fecal level of taurine (taurine: 0.543, taurine+Na: 0.909), a by-product of endogenous oxidative cysteine metabolism, in HuGF-*Il10*^-/-^ mice compared to vehicle group (**Fig. 6D**). Studies have shown that taurine has protective effect on colitis^41–43^. We also observed decreased fecal level of histamine (Histamine: 0.463) and its derivate (N-methylhistamine) after AuPhos treatment (**Fig. 6D**). AuPhos modulates metabolites involved in all three components of energy metabolisms; viz. energy substrate transfer, OxPhos and high-energy phosphate bond transfer and utilization (**Fig. 7A**). During IBD, inflammation-induced mitochondrial dysfunction leads to incomplete FAO that imbalance carnitine shuttle system, thereby decreasing carnitine levels and increasing un-utilized acylcarnitines88-91. Interestingly, AuPhos treatment increased the level of L-carnitine (L-Carnitine+Cl: 0.307, L-Carnitine: 0.872) and decreased acylcarnitines (Acetylcarnitine: 0.515, Isovalerylcarnitine: 0.840, Isovalerylcarnitine, Deoxycarnitine: 0.890, O-Decanoyl-L-carnitine, O-Propanoylcarnitine) in fecal samples from HuGF-Il10-/- mice (**Fig. 7B**). We also observed significantly increased levels of TCA intermediates (2-Hydroxyglutarate+water: 0.805, 2-Hydroxyglutarate+Na: 0.746, 2-Oxoglutarate, succinate-H2O: 0.769, Succinate+Na: 0.793, Succinate: 0.842) (**Fig. 7C**) and creatine (indicating increased ATP production in IECs) in the fecal samples from AuPhos treated HuGF-Il10-/- mice (**Fig. 7D**). These findings indicate that AuPhos-enhanced IEC mitochondrial function reduces enteric O2 delivery, which corrects disease-associated metabolomics by restoring short-chain fatty acids, SBA, AA and IEC energy metabolism.

**Figure 6.**
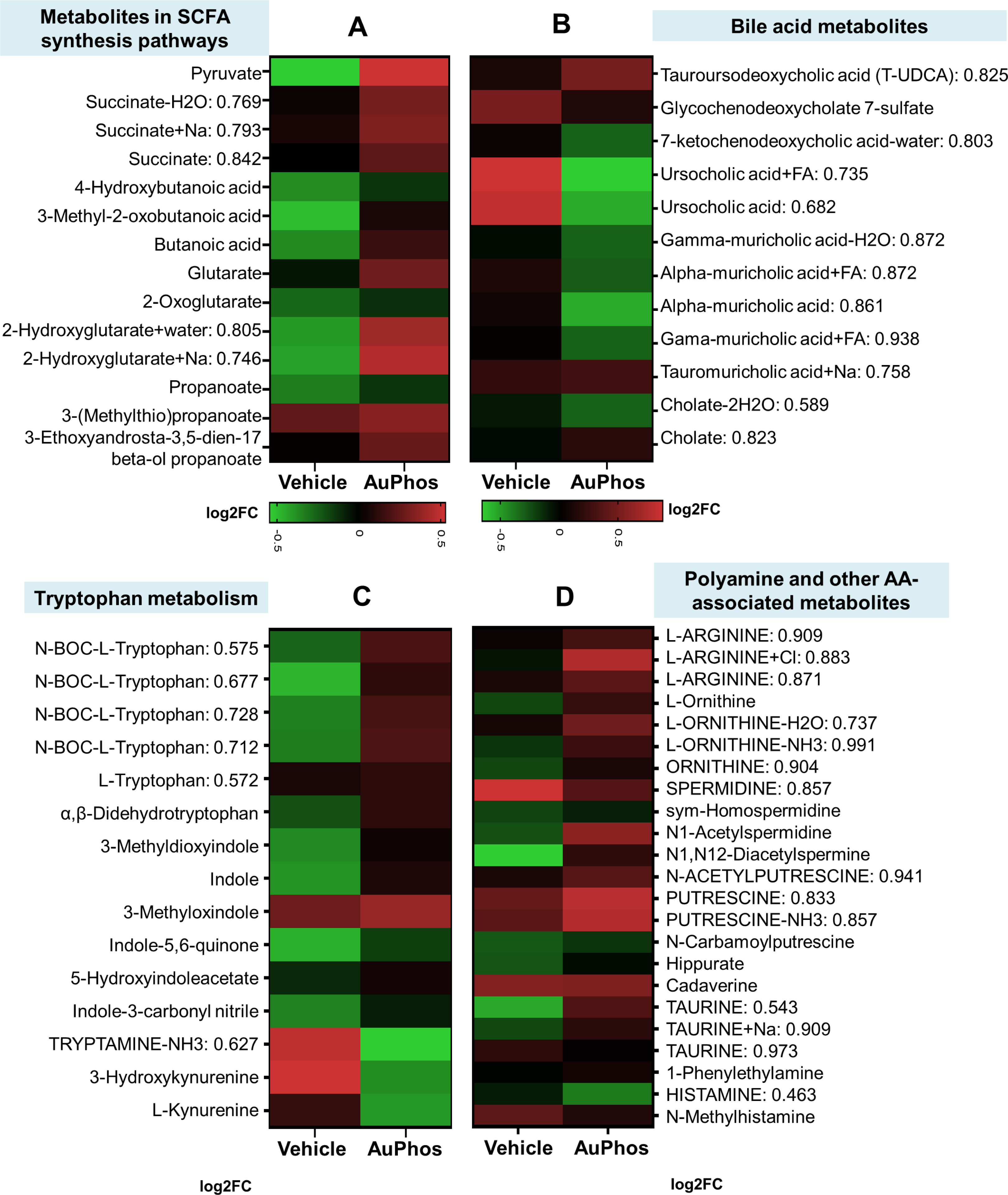
AuPhos effect on key metabolites associated with SCFA synthesis and bile acid metabolism in fecal samples from Hu-*Il10-/-* mice. Heat map showing metabolites upregulated (red) or downregulated (green) in AuPhos treated mice (n=6) compared to vehicle group (n=7). These data indicate that **AuPhos** increases SCFA synthesis and deconjugation of PBA into SBAs. AuPhos effect on key metabolites associated with AA and polyamine metabolism in fecal samples from Hu-*Il10-/-* mice. Heat map showing metabolites upregulated (red) or downregulated (green) in AuPhos treated mice (n=6) compared to vehicle group (n=7). These data indicate that **AuPhos** increases tryptophan metabolism and other AAs involved in polyamine biosynthesis, and reduction in inflammatory histamine production.

**Figure 7.**
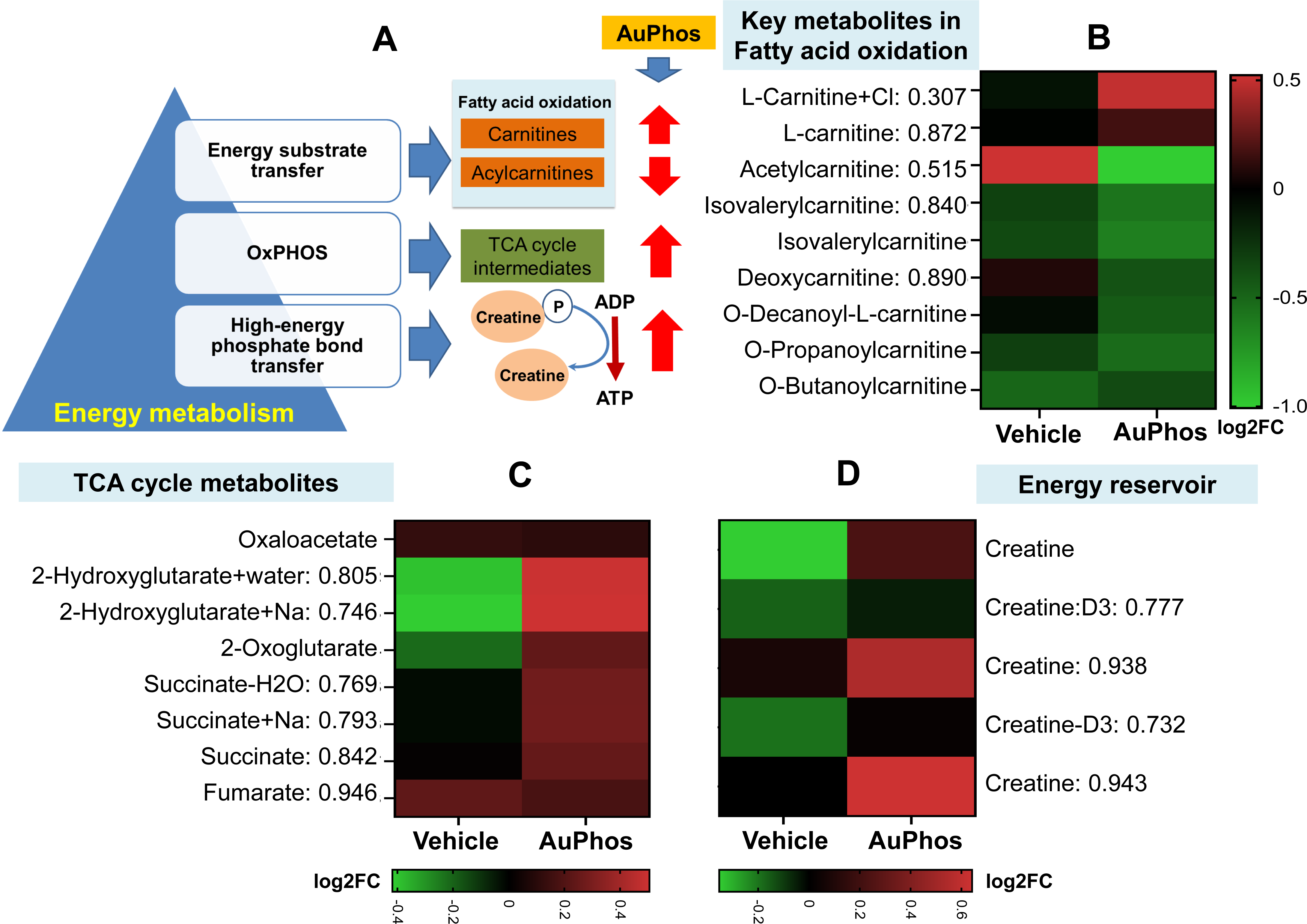
AuPhos effect on key metabolites associated with energy metabolism in fecal samples from Hu-*Il10-/-* mice. Heat map showing metabolites upregulated (red) or downregulated (green) in AuPhos treated mice (n=6) compared to vehicle group (n=7). These data indicate that **AuPhos** increases FAO, TCA metabolites and an energy reservoir Creatine.

## DISCUSSION

In healthy individuals, intestines are lined with differentiated intestinal epithelial cells (IECs) that have relatively high levels of mitochondria that consume oxygen (O2) by oxidative phosphorylation (OXPHOS) creating a hypoxic microenvironment (**Fig 5**). O2 consumption reduces O2 availability to the microbiome and promotes a healthy anaerobic environment dominated by obligate anaerobes (OAs). These OAs ferment dietary fibers and produce short chain fatty acids (SCFA) that in turn provide substrate for IEC OXPHOS. During IBD, inflammation-induced oxidant damage impairs mitochondrial function thereby limiting OXPHOS and converting IECs to a greater glycolytic metabolism that consumes less O2^34^. Reduced IEC O2 consumption increases the amount of O2 emanating from the colonic epithelium. Increased enteric O2 promotes blooms of facultative anaerobes (e.g. *Proteobacteria*) and restricts obligate anaerobes (e.g. *Firmicutes*)^2,34^. Dysbiotic metabolites negatively affect host metabolism and immunity^44^. Given the potential that **AuPhos** upregulates IEC mitochondrial function^28,29^, attenuates colitis and corrects dysbiosis in humanized *Il10^-/-^* mice (**Fig 5**), we investigated the effect of AuPhos in rewiring the IBD-associated dysbiotic metabolism using untargeted metabolomics on stool samples collected from **AuPhos**- or vehicle-treated Hu-10-/- and WT mice (**Fig 6,7**). AuPhos effects on microbial metabolites was determined in stool samples from germ-free (GF) 129.SvEv WT or *Il10^-/-^* mice reconstituted with human IBD stool (Hu-*Il10-/-*). Mice were treated orally with AuPhos (10-mg/kg; q2d; 2wk) or vehicle. Accumulating evidences suggest that fecal samples from IBD patients have decreased level of SCFAs (butyrate, propionate and acetate) in varying degrees^45,46^. SCFAs play a crucial role in IEC energy metabolism, supporting IEC proliferation and gut barrier integrity, inhibiting pro-inflammatory cytokines, and modulating systemic immune response through host receptor signals^47^. Microbiota-derived butyrate particularly suppresses intestinal inflammation by activating G-protein receptor 41 (GPR-41) and inhibiting histone deacetylase (HDAC) to IL-22 production^48^. AuPhos significantly altered the metabolites associated with butyrate synthesis in Hu-*Il10-/-* mice (**Fig 6A**). AuPhos significantly increased pyruvate, succinate, 4-hydroxybutyrate, 2-hydroxyglutarate, and butanoic acid (butyrate) compared to vehicle treated mice, suggesting an upregulation of pyruvate, 4-aminobutyrate (4Ab) and glutarate pathways of butyrate production (**Fig 6A**). AuPhos also marginally increased the level of propionate and its derivates (**Fig 6A**). Another important feature of commensal gut bacteria is conversion of primary BAs (PBAs) to secondary BAs (SBAs), which is mediated by the enzymes associated with dihydroxylation and deconjugation^36^. Studies have shown illustrate the important role of SBAs in ameliorating intestinal inflammation and promoting colonic epithelial restitution by decreasing pro-inflammatory cytokines and inhibiting IEC apoptosis^49^. Earlier MS-based metabolomic studies show that BA metabolism is perturbed in IBD, with elevated PBAs and reduced level of SBAs^50^. Significant reduction in fecal SBAs has been shown to be positively correlated with *Roseburia, Clostridium IV, Butyricicoccus, and Faecalibacterium* in UC patients^50^. AuPhos reduced IBD-associated PBAs (Ketochenodeoxycholic, Glycochenodeoxycholic Ursocholic acid, Cholate, Muricholic acid, and different derivatives of these PBAs) with concomitant increase in SBA (Tauroursodeoxycholic acid, TUDCA) (**Fig 6B**). Pre-clinical studies on experimental mice (TNFΔ^ARE/WT^) have shown that exogenous supplementation of TUDCA protects bile acid homeostasis under inflammatory conditions and suppresses CD-like ileitis^51^.

Intestinal inflammation also significantly alters amino acid metabolism that correlates with disease severity^37^. Additionally, metagenomic analysis of gut microbiome of IBD patients have revealed downregulation of amino acid (AA) biosynthesis genes and upregulation of AA transporter genes, suggesting reduced AA synthesis and concomitant increase in AA utilization by dysbiotic microbiota in inflamed gut^52,53^. Besides, inflammation also induces differential utilization of host immune cells. For instance, certain AAs modulate inflammation by reciprocal regulation of macrophages and some are critical in promoting T-cell proliferation and differentiation^54,55^. Thus, inflammation may induce metabolic reprogramming that increases demand for certain AA by host cells and the gut microbiota. Tryptophan (TRP) metabolism in particular plays an important role in intestinal barrier function, gut hormone secretion, gut motility and systemic immune response by inhibiting the release of pro-inflammatory factors by macrophages^56–58^ after binding to 5-hydroxytryptamine receptor (5-HTR) in macrophage leading to serotonin (5-HT) production. Accumulating evidence suggests that microbiota-derived TRP metabolites play a crucial role in mediating host-microbial cross-talk by binding to cytosolic aryl hydrocarbon receptor or pregnane X receptor^59^. Targeted or untargeted metabolomics have revealed that TRP metabolism is perturbed and indoles and derivatives are significantly depleted in IBD patients^60–62^. Studies have shown that administration of TRP and its metabolites (indole, indole-3-aldehyde, Indole-3-propionic acid and indole-3-acetic acid) reversed colitis-associated microbial dysbiosis, colonic inflammation and restores intestinal barrier integrity^63–66^. Furthermore, dietary TRP deficiency was associated with the exacerbation of colitis, and indole-3-carbinol treatment effectively corrects the TNBS-induced dysbiotic microbial composition, selectively increasing the abundance of *Roseburia*, which is an excellent butyrate producer^65^. Reports show strong association of tryptophan deficiency with dysbiosis of the gut microbiota leading to gastrointestinal and systemic inflammation^67,68^. Our untargeted metabolomic analysis show that AuPhos treatment significantly increased levels of fecal TRP (N-BOC-L-Tryptophan: 0.575, N-BOC-L-Tryptophan: 0.677, N-BOC-L-Tryptophan: 0.728, N-BOC-L-Tryptophan: 0.712, L-Tryptophan: 0.572, α,β-Didehydrotryptophan) and various indole-derivatives (3-Methyldioxyindole, Indole, 3-Methyloxindole, 5-Hydroxyindoleacetate, Indole-3-carbonyl nitrile) in Hu-*Il10-/-* mice compared to vehicle control (**Fig 6C**). Further, kynurenine (KYN) pathway is the major catabolic pathway of TRP accounting for 95% of its degradation by indoleamine 2,3-dioxygenase 1 (IDO1), activated under inflammatory environment^69^. There is a strong connection between increased KYN levels and IBD-associated depression^70^ Colonic biopsies of IBD patients show elevated levels of the IDO messenger RNA and KYN/TRP^71^, primarily due to proinflammatory cytokines (IFN-γ, IL-1, and IL-6)-mediated activation of IDO^72^. Several studies show that KYN/TRP ratio is associated with important biomarkers of disease activity, C-reactive protein, sedimentation rate^73^, and endoscopic score^74^. Interestingly, phylum *Proteobacteria*-derived lipopolysaccharide (LPS) stimulated colonic immune responses upregulates the indoleamine 2,3-dioxygenase 1 (IDO1)-mediated Kyn pathway, leading to TRP depletion and KYN accumulation in the circulation^75^. AuPhos treatment reduced the level of KYN (L-Kynurenine) and its hydroxy-derivatives (3-Hydroxykynurenine) in Hu-*Il10-/-* mice compared to vehicle treated group (**Fig 6C**). Colonic microbiota ferment proteins leading to the production of certain bioactive end-products known as polyamines, with L-arginine being the primary precursor. Putrescine is the most abundant polyamine prevalent in large intestine of humans followed by spermine, spermidine and cadaverine^76–78^. Arginine is converted into putrescine by intermediate production of either L-ornithine or N-Carbamoyl-putrescine^79^. Putrescine could further be converted into spermidine and spermine. Earlier studies demonstrate the importance of polyamines in accelerating the proliferation and wound healing process in the intestinal epithelium^80,81^. Microbial-derived putrescine reinforce colonic epithelial proliferation, IEC OXPHOS, increases anti-inflammatory M2 macrophages in the colon ameliorates DSS-induced colitis^82,83^. Polyamines enhance hypunisation of eukaryotic initiation factor (EIF5A), which promotes efficient expression of mitochondrial OXPHOS proteins, thereby increasing O2 consumption, mitochondrial function and help maintain mucosal homeostasis in the intestine^82–84^. Studies have shown that putrescine also serves as a direct energy source for enterocytes by converting into succinate and fuels OXPHOS^85^. **AuPhos** treatment significantly increased the fecal levels of arginine (L-Arginine: 0.909, L-Arginine+Cl: 0.883, L-Arginine: 0.871), ornithine (L-Ornithine, L-Ornithine-H2O: 0.737, L-Ornithine-NH3: 0.991, Ornithine: 0.904), N-Carbamoylputrescine and putrescine (N-Acetylputrescine: 0.941, Putrescine: 0.833, Putrescine-NH3: 0.857) compared to vehicle group in Hu-*Il10-/-* mice (**Fig. 6D**). Our metabolomic data showed increased fecal level of double acetylated forms of spermine (N1,N12-Diacetylspermine), which is a catabolic product of spermine for inter-conversion back to spermidine or putrescine^38–40^. Under high mitochondrial activity (with active ATP production), spermine is transported back and forth over the mitochondria membrane in parallel with ADP and phosphate via adenine nucleotide translocase (ANT)^86^. This process is believed to have a restorative effect on oxidative functions of mitochondria that further upregulated its ATP production^87,88^. Studies also show that spermine regulate calcium transport into mitochondria, which controls pyruvate influx (via pyruvate dehydrogenase complex) and its conversion acetyl-CoA, thereby increased mitochondrial metabolic rate^89,90^. Besides putrescine and spermine, **AuPhos** did not show much effect on level of spermidine and cadaverine in Hu-*Il10-/-* mice compared to vehicle treatment **(Fig. 6D).** Our untargeted metabolomics showed increased fecal level of taurine (Taurine: 0.543, Taurine+Na: 0.909), a by-product of endogenous oxidative cysteine metabolism, in Hu-*Il10-/-* mice compared to vehicle group (**Fig. 6D**). Studies show that taurine has protective effect on colitis^41–43^, It activates NLRP6-IL-18-AMPs pathway facilitating reduction in colitis severity and reversal of dysbiotic microbiota^91^. Besides, high level of taurine in the gut associates with increased intestinal epithelial integrity by enhancing tight junctions, thereby reduced leaky gut and inflammation^92^. We observed decreased fecal level of Histamine (Histamine: 0.463) and its derivate (N-Methylhistamine) after **AuPhos** treatment (**Fig. 6D**). Histamine is an oxidative decarboxylation product of L-Histidine mediated by histidine decarboxylase (HDC) enzyme^93^, which is also present in large number of gut microbiota^94–96^. Dysbiotic gut has significantly higher abundance of histamine-secreting (e.g. *Enterobacteriaceae*) and histamine-intolerant bacteria^97–99^. Studies show increased mucosal histamine levels in patients with IBD, with increased urine levels of N-methylhistamine correlated with disease activity^100–102^. Expression and functional activity of histamine receptor (H2R) is altered in IBD patients, a notion supported by the study where H2R blockage resulted in more severe inflammatory disease in the murine T-cell transfer colitis model^103^. Histamine N-methyltransferase (HNMT) and diaminoxidase (DAO) are the two enzymes responsible for histamine degradation and tolerance, respectively^104,105^. Studies show that DAO polymorphisms and reduced HNMT gene expression in inflamed mucosa are associated with an increased risk of IBD^103,106,107^. Histamine also drives severity of innate inflammation via histamine 4 receptor in murine experimental colitis^108^. **AuPhos** also modulates metabolites involved in all three components of energy metabolisms; viz. energy substrate transfer, OXPHOS and high-energy phosphate bond transfer and utilization (**Fig. 7**). Carnitines and acylcarnitines are the primary metabolites reflective of mitochondrial metabolism that facilitate fatty-acid beta-oxidation (FAO) in the mitochondrial matrix^109,110^. Carnitine is an important component of shuttle system that imports long-chain fatty-acids (LCFAs) inside the mitochondria for subsequent FAO^109,111,112^. LCFAs are converted into acyl-CoA, which then combines with free carnitines to produce acylcarnitines with the help of enzyme carnitine palmitoyltransferase-1. These acylcarnitines are then shuttled across inner mitochondrial membrane into the matrix, where they get converted back into acyl-CoA and free carnitine. Free carnitines are exported out and acyl-CoA produces acetyl-CoA via β-oxidation to serve as an energy substrate for TCA cycle. During IBD, inflammation-induced mitochondrial dysfunction leads to incomplete FAO that imbalance carnitine shuttle system, thereby decreasing carnitine levels and increasing un-utilized acylcarnitines^113–116^. Studies show that polymorphisms in *SLC22A5* gene that encodes for the carnitine transporter OCTN2, is an associated risk factor for IBD^117,118^. Interestingly, **AuPhos** treatment increases L-carnitine (L-Carnitine+Cl: 0.307, L-Carnitine: 0.872) and decreases acylcarnitines (Acetylcarnitine: 0.515, Isovalerylcarnitine: 0.840,

Isovalerylcarnitine, Deoxycarnitine: 0.890, O-Decanoyl-L-carnitine, O-Propanoylcarnitine) in fecal samples from Hu-*Il10-/-* mice compared to vehicle treatment (**Fig. 7B**). Recent studies show that increased fecal level of intestinal acylcarnitines promotes the growth of *Enterobacteriaceae* and is considered as a biomarker of dysbiosis in IBD^119^. Besides FAO disruption, multiple studies show altered energy metabolism involving significant decrease in serum, plasma, urine and tissue levels of TCA intermediates (citrate, aconitate, α-ketoglutarate, succinate, fumarate, and malate) in IBD compared to healthy controls^120–124^. However, no change was observed in fecal samples. We observed significantly increased levels of TCA intermediates (2-Hydroxyglutarate+water: 0.805, 2-Hydroxyglutarate+Na: 0.746, 2-Oxoglutarate, Succinate-H2O: 0.769, Succinate+Na: 0.793, Succinate: 0.842, Malate) and creatine (indicating increased ATP production in IECs) in the fecal samples from **AuPhos-**treated Hu-*Il10-/-* mice compared to vehicle control. We believe that these alterations in Kreb’s cycle metabolites are due to shedding of **AuPhos**-treated mucosal IEC and permeation of metabolites into the microbiome. These findings indicate that AuPhos-enhanced IEC mitochondrial function reduces enteric O2 delivery, which corrects disease-associated metabolomics by restoring SCFA production, SBA, AA and IEC energy metabolism.

Together these exciting new data indicate that regulation of mucosal mitochondrial function offers a unique opportunity for correcting mucosal inflammatory responses as well as correcting dysbiotic microbiomes. We post that targeting this pathway in IBD (and other disease states) provide an opportunity for maintaining effects of biologic agents and prevent disease recurrence.

## Acknowledgement

We acknowledge the support of university of Kentucky (UKY) and Veteran Affairs (VA) animal facilities. Human biopsy collection was approved by the UKY IRB. This work was supported in part by a VA Merit Award [1I01CX001353-01A1], Awuah grant [NIH RO1CA258421-01; NIH COBRE P20 GM130456-01A1], Barrett RO1 grant [2R01 DK095662-10], Barrett Litwin Award (993820) and patents [PCT/US21/43774; PCT/US21/52719].

---

**Supplementary Fig. 1:**
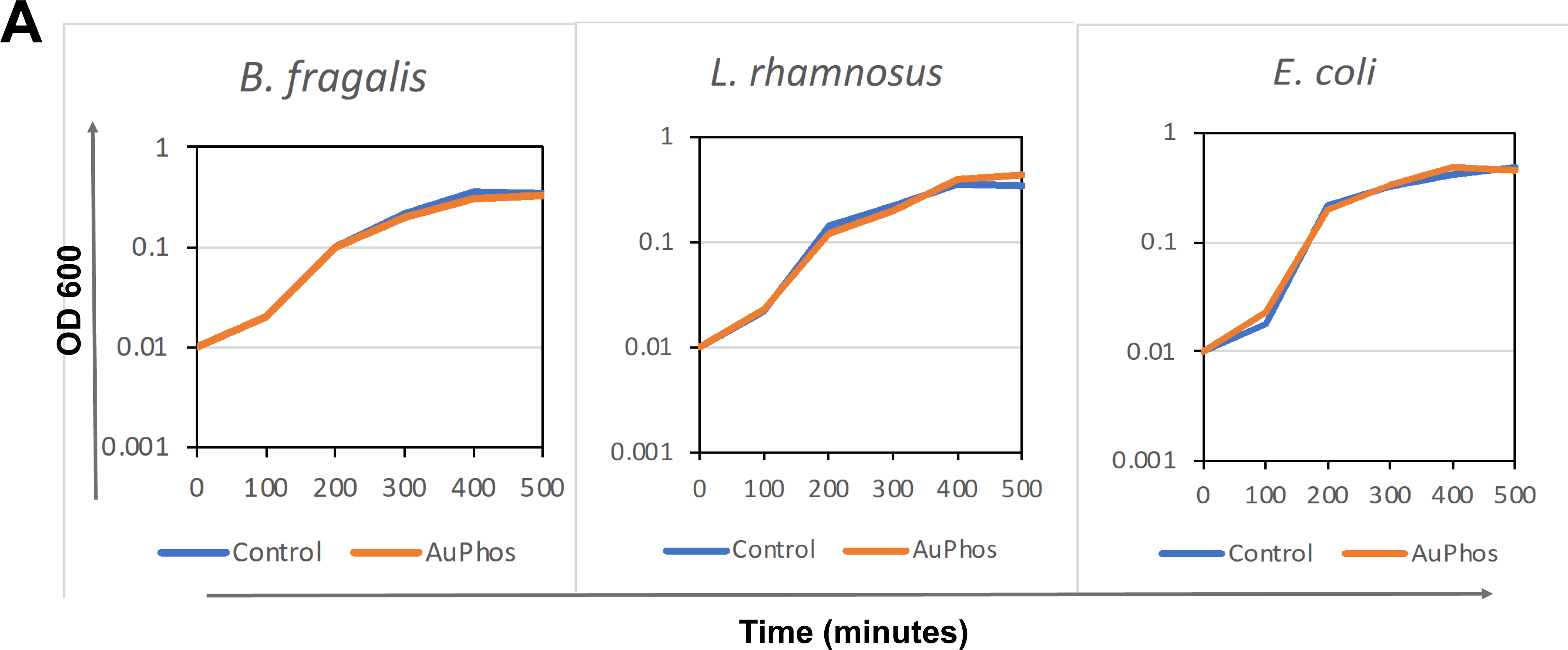
Absence of AuPhos direct effects on bacteria.

## REFERENCES

1 Rath, E. & Haller, D. Intestinal epithelial cell metabolism at the interface of microbial dysbiosis and tissue injury. Mucosal Immunol 15, 595–604 (2022). 10.1038/s41385-022-00514-x

2 Litvak, Y., Byndloss, M. X., Tsolis, R. M. & Baumler, A. J. Dysbiotic Proteobacteria expansion: a microbial signature of epithelial dysfunction. Curr Opin Microbiol 39, 1–6 (2017). 10.1016/j.mib.2017.07.003

3 Litvak, Y., Byndloss, M. X. & Baumler, A. J. Colonocyte metabolism shapes the gut microbiota. Science 362 (2018). 10.1126/science.aat9076

4 Lloyd-Price, J. et al. Multi-omics of the gut microbial ecosystem in inflammatory bowel diseases. Nature 569, 655–662 (2019). 10.1038/s41586-019-1237-9

5 Khan, I. et al. Alteration of Gut Microbiota in Inflammatory Bowel Disease (IBD): Cause or Consequence? IBD Treatment Targeting the Gut Microbiome. Pathogens 8 (2019). 10.3390/pathogens8030126

6 Qiu, X., Zhang, M., Yang, X., Hong, N. & Yu, C. Faecalibacterium prausnitzii upregulates regulatory T cells and anti-inflammatory cytokines in treating TNBS-induced colitis. J Crohns Colitis 7, e558–568 (2013). 10.1016/j.crohns.2013.04.002

7 Gaboriau-Routhiau, V. et al. The key role of segmented filamentous bacteria in the coordinated maturation of gut helper T cell responses. Immunity 31, 677–689 (2009). 10.1016/j.immuni.2009.08.020

8 Ivanov, II et al. Induction of intestinal Th17 cells by segmented filamentous bacteria. Cell 139, 485–498 (2009). 10.1016/j.cell.2009.09.033

9 Zeng, M. Y., Inohara, N. & Nunez, G. Mechanisms of inflammation-driven bacterial dysbiosis in the gut. Mucosal Immunol 10, 18–26 (2017). 10.1038/mi.2016.75

10 Jans, D. & Cleynen, I. The genetics of non-monogenic IBD. Hum Genet 142, 669–682 (2023). 10.1007/s00439-023-02521-9

11 Ho, G. T. et al. MDR1 deficiency impairs mitochondrial homeostasis and promotes intestinal inflammation. Mucosal Immunol 11, 120–130 (2018). 10.1038/mi.2017.31

12 Saxena, A., Lopes, F., Poon, K. K. H. & McKay, D. M. Absence of the NOD2 protein renders epithelia more susceptible to barrier dysfunction due to mitochondrial dysfunction. Am J Physiol Gastrointest Liver Physiol 313, G26–G38 (2017). 10.1152/ajpgi.00070.2017

13 Haberman, Y. et al. Ulcerative colitis mucosal transcriptomes reveal mitochondriopathy and personalized mechanisms underlying disease severity and treatment response. Nat Commun 10, 38 (2019). 10.1038/s41467-018-07841-3

14 Kugathasan, S. et al. Prediction of complicated disease course for children newly diagnosed with Crohn’s disease: a multicentre inception cohort study. Lancet 389, 1710–1718 (2017). 10.1016/S0140-6736(17)30317-3

15 Hsieh, S. Y. et al. Comparative proteomic studies on the pathogenesis of human ulcerative colitis. Proteomics 6, 5322–5331 (2006). 10.1002/pmic.200500541

16 Mottawea, W. et al. Altered intestinal microbiota-host mitochondria crosstalk in new onset Crohn’s disease. Nat Commun 7, 13419 (2016). 10.1038/ncomms13419

17 Alula, K. M. et al. Targeting Mitochondrial Damage as a Therapeutic for Ileal Crohn’s Disease. Cells 10 (2021). 10.3390/cells10061349

18 Sifroni, K. G. et al. Mitochondrial respiratory chain in the colonic mucosal of patients with ulcerative colitis. Mol Cell Biochem 342, 111–115 (2010). 10.1007/s11010-010-0474-x

19 Santhanam, S. et al. Mitochondrial electron transport chain complex dysfunction in the colonic mucosa in ulcerative colitis. Inflamm Bowel Dis 18, 2158–2168 (2012). 10.1002/ibd.22926

20 Santhanam, S., Venkatraman, A. & Ramakrishna, B. S. Impairment of mitochondrial acetoacetyl CoA thiolase activity in the colonic mucosa of patients with ulcerative colitis. Gut 56, 1543–1549 (2007). 10.1136/gut.2006.108449

21 Byndloss, M. X. et al. Microbiota-activated PPAR-gamma signaling inhibits dysbiotic Enterobacteriaceae expansion. Science 357, 570–575 (2017). 10.1126/science.aam9949

22 Spiga, L. et al. An Oxidative Central Metabolism Enables Salmonella to Utilize Microbiota-Derived Succinate. Cell Host Microbe 22, 291–301 e296 (2017). 10.1016/j.chom.2017.07.018

23 El Kaoutari, A., Armougom, F., Gordon, J. I., Raoult, D. & Henrissat, B. The abundance and variety of carbohydrate-active enzymes in the human gut microbiota. Nat Rev Microbiol 11, 497–504 (2013). 10.1038/nrmicro3050

24 Krautkramer, K. A., Fan, J. & Backhed, F. Gut microbial metabolites as multi-kingdom intermediates. Nat Rev Microbiol 19, 77–94 (2021). 10.1038/s41579-020-0438-4

25 Louis, P. & Flint, H. J. Formation of propionate and butyrate by the human colonic microbiota. Environ Microbiol 19, 29–41 (2017). 10.1111/1462-2920.13589

26 Rousta, E. et al. The Emulsifier Carboxymethylcellulose Induces More Aggressive Colitis in Humanized Mice with Inflammatory Bowel Disease Microbiota Than Polysorbate-80. Nutrients 13 (2021). 10.3390/nu13103565

27 Kim, J. H. et al. Anticancer gold(iii)-bisphosphine complex alters the mitochondrial electron transport chain to induce in vivo tumor inhibition. Chem Sci 12, 7467–7479 (2021). 10.1039/d1sc01418h

28 Kapur, N. et al. Sa1533: NOVEL ORAL THERAPY THAT INCREASES MITOCHONDRIAL FUNCTION AND ATTENUATES COLITIS. Gastroenterology 162, S-405 (2022). 10.1016/S0016-5085(22)60960-0

29 Kapur, N. et al. Sa1534: NOVEL GOLD COMPOUND THAT REGULATES CELL METABOLISM TO PROMOTE CRYPT FISSIONING IN IBD. Gastroenterology 162, S-405 (2022). 10.1016/S0016-5085(22)60961-2

30 Mohamed, R. et al. AUPHOS, A NOVEL THERAPEUTIC THAT IMPROVES MITOCHONDRIAL FUNCTION AND AMELIORATES CHRONIC COLITIS. Gastroenterology 162, S2–S3 (2022). 10.1053/j.gastro.2021.12.011

31 Wang, H. et al. Chronic Model of Inflammatory Bowel Disease in IL-10(-/-) Transgenic Mice: Evaluation with Ultrasound Molecular Imaging. Theranostics 9, 6031–6046 (2019). 10.7150/thno.37397

32 Brown, J. B., Lee, G., Grimm, G. R. & Barrett, T. A. Therapeutic benefit of pentostatin in severe IL-10-/- colitis. Inflamm Bowel Dis 14, 880–887 (2008). 10.1002/ibd.20410

33 Berg, D. J. et al. Rapid development of colitis in NSAID-treated IL-10-deficient mice. Gastroenterology 123, 1527–1542 (2002). 10.1053/gast.2002.1231527

34 Rivera-Chavez, F., Lopez, C. A. & Baumler, A. J. Oxygen as a driver of gut dysbiosis. Free Radic Biol Med 105, 93–101 (2017). 10.1016/j.freeradbiomed.2016.09.022

35 Bhogoju, S. et al. A NOVEL DRUG THERAPY INDUCES MITOCHONDRIAL BIOGENESIS AND ATTENUATES COLITIS. Gastroenterology 164, S70 (2023). 10.1053/j.gastro.2023.03.134

36 Ridlon, J. M., Kang, D. J. & Hylemon, P. B. Bile salt biotransformations by human intestinal bacteria. J Lipid Res 47, 241–259 (2006). 10.1194/jlr.R500013-JLR200

37 Sugihara, K., Morhardt, T. L. & Kamada, N. The Role of Dietary Nutrients in Inflammatory Bowel Disease. Front Immunol 9, 3183 (2018). 10.3389/fimmu.2018.03183

38 Bae, D. H., Lane, D. J. R., Jansson, P. J. & Richardson, D. R. The old and new biochemistry of polyamines. Biochim Biophys Acta Gen Subj 1862, 2053–2068 (2018). 10.1016/j.bbagen.2018.06.004

39 Pegg, A. E. Mammalian polyamine metabolism and function. IUBMB Life 61, 880–894 (2009). 10.1002/iub.230

40 Casero, R. A., Jr. & Pegg, A. E. Spermidine/spermine N1-acetyltransferase--the turning point in polyamine metabolism. FASEB J 7, 653–661 (1993).

41 Son, M., Ko, J. I., Kim, W. B., Kang, H. K. & Kim, B. K. Taurine can ameliorate inflammatory bowel disease in rats. Adv Exp Med Biol 442, 291–298 (1998). 10.1007/978-1-4899-0117-0_37

42 Son, M. W. et al. Protective effect of taurine on TNBS-induced inflammatory bowel disease in rats. Arch Pharm Res 21, 531–536 (1998). 10.1007/BF02975370

43 Zhao, Z. et al. Attenuation by dietary taurine of dextran sulfate sodium-induced colitis in mice and of THP-1-induced damage to intestinal Caco-2 cell monolayers. Amino Acids 35, 217–224 (2008). 10.1007/s00726-007-0562-8

44 Konturek, P. C. [Gut microbiota and chronic inflammatory bowel disease]. MMW Fortschr Med 164, 12–15 (2022). 10.1007/s15006-022-1230-3

45 Marchesi, J. R. et al. Rapid and noninvasive metabonomic characterization of inflammatory bowel disease. J Proteome Res 6, 546–551 (2007). 10.1021/pr060470d

46 Machiels, K. et al. A decrease of the butyrate-producing species Roseburia hominis and Faecalibacterium prausnitzii defines dysbiosis in patients with ulcerative colitis. Gut 63, 1275–1283 (2014). 10.1136/gutjnl-2013-304833

47 Deleu, S., Machiels, K., Raes, J., Verbeke, K. & Vermeire, S. Short chain fatty acids and its producing organisms: An overlooked therapy for IBD? EBioMedicine 66, 103293 (2021). 10.1016/j.ebiom.2021.103293

48 Yang, W. et al. Intestinal microbiota-derived short-chain fatty acids regulation of immune cell IL-22 production and gut immunity. Nat Commun 11, 4457 (2020). 10.1038/s41467-020-18262-6

49 Ward, J. B. J. et al. Ursodeoxycholic acid and lithocholic acid exert anti-inflammatory actions in the colon. Am J Physiol Gastrointest Liver Physiol 312, G550–G558 (2017). 10.1152/ajpgi.00256.2016

50 Yang, Z. H. et al. Altered profiles of fecal bile acids correlate with gut microbiota and inflammatory responses in patients with ulcerative colitis. World J Gastroenterol 27, 3609–3629 (2021). 10.3748/wjg.v27.i24.3609

51 Van den Bossche, L., et al. Tauroursodeoxycholic acid protects bile acid homeostasis under inflammatory conditions and dampens Crohn’s disease-like ileitis. Lab Invest 97, 519–529 (2017). 10.1038/labinvest.2017.6

52 Morgan, X. C. et al. Dysfunction of the intestinal microbiome in inflammatory bowel disease and treatment. Genome Biol 13, R79 (2012). 10.1186/gb-2012-13-9-r79

53 Davenport, M. et al. Metabolic alterations to the mucosal microbiota in inflammatory bowel disease. Inflamm Bowel Dis 20, 723–731 (2014). 10.1097/MIB.0000000000000011

54 Ren, W. et al. Amino-acid transporters in T-cell activation and differentiation. Cell Death Dis 8, e2655 (2017). 10.1038/cddis.2016.222

55 Langston, P. K., Shibata, M. & Horng, T. Metabolism Supports Macrophage Activation. Front Immunol 8, 61 (2017). 10.3389/fimmu.2017.00061

56 Haq, S., Grondin, J. A. & Khan, W. I. Tryptophan-derived serotonin-kynurenine balance in immune activation and intestinal inflammation. FASEB J 35, e21888 (2021). 10.1096/fj.202100702R

57 de las Casas-Engel, M., et al. Serotonin skews human macrophage polarization through HTR2B and HTR7. J Immunol 190, 2301–2310 (2013). 10.4049/jimmunol.1201133

58 Mawe, G. M. & Hoffman, J. M. Serotonin signalling in the gut--functions, dysfunctions and therapeutic targets. Nat Rev Gastroenterol Hepatol 10, 473–486 (2013). 10.1038/nrgastro.2013.105

59 Stockinger, B., Shah, K. & Wincent, E. AHR in the intestinal microenvironment: safeguarding barrier function. Nat Rev Gastroenterol Hepatol 18, 559–570 (2021). 10.1038/s41575-021-00430-8

60 Franzosa, E. A. et al. Author Correction: Gut microbiome structure and metabolic activity in inflammatory bowel disease. Nat Microbiol 4, 898 (2019). 10.1038/s41564-019-0442-5

61 Nikolaus, S. et al. Increased Tryptophan Metabolism Is Associated With Activity of Inflammatory Bowel Diseases. Gastroenterology 153, 1504–1516 e1502 (2017). 10.1053/j.gastro.2017.08.028

62 Lamas, B. et al. CARD9 impacts colitis by altering gut microbiota metabolism of tryptophan into aryl hydrocarbon receptor ligands. Nat Med 22, 598–605 (2016). 10.1038/nm.4102

63 Scott, S. A., Fu, J. & Chang, P. V. Microbial tryptophan metabolites regulate gut barrier function via the aryl hydrocarbon receptor. Proc Natl Acad Sci U S A 117, 19376–19387 (2020). 10.1073/pnas.2000047117

64 Cervantes-Barragan, L. et al. Lactobacillus reuteri induces gut intraepithelial CD4(+)CD8alphaalpha(+) T cells. Science 357, 806–810 (2017). 10.1126/science.aah5825

65 Busbee, P. B. et al. Indole-3-carbinol prevents colitis and associated microbial dysbiosis in an IL-22-dependent manner. JCI Insight 5 (2020). 10.1172/jci.insight.127551

66 Zelante, T. et al. Tryptophan catabolites from microbiota engage aryl hydrocarbon receptor and balance mucosal reactivity via interleukin-22. Immunity 39, 372–385 (2013). 10.1016/j.immuni.2013.08.003

67 Yusufu, I. et al. A Tryptophan-Deficient Diet Induces Gut Microbiota Dysbiosis and Increases Systemic Inflammation in Aged Mice. Int J Mol Sci 22 (2021). 10.3390/ijms22095005

68 He, F. et al. Functions and Signaling Pathways of Amino Acids in Intestinal Inflammation. Biomed Res Int 2018, 9171905 (2018). 10.1155/2018/9171905

69 Badawy, A. A. Kynurenine Pathway of Tryptophan Metabolism: Regulatory and Functional Aspects. Int J Tryptophan Res 10, 1178646917691938 (2017). 10.1177/1178646917691938

70 Chen, L. M. et al. Tryptophan-kynurenine metabolism: a link between the gut and brain for depression in inflammatory bowel disease. J Neuroinflammation 18, 135 (2021). 10.1186/s12974-021-02175-2

71 Wolf, A. M. et al. Overexpression of indoleamine 2,3-dioxygenase in human inflammatory bowel disease. Clin Immunol 113, 47–55 (2004). 10.1016/j.clim.2004.05.004

72 Ciorba, M. A. Indoleamine 2,3 dioxygenase in intestinal disease. Curr Opin Gastroenterol 29, 146–152 (2013). 10.1097/MOG.0b013e32835c9cb3

73 Gupta, N. K. et al. Serum analysis of tryptophan catabolism pathway: correlation with Crohn’s disease activity. Inflamm Bowel Dis 18, 1214–1220 (2012). 10.1002/ibd.21849

74 Sofia, M. A. et al. Tryptophan Metabolism through the Kynurenine Pathway is Associated with Endoscopic Inflammation in Ulcerative Colitis. Inflamm Bowel Dis 24, 1471–1480 (2018). 10.1093/ibd/izy103

75 Sun, P. et al. High-fat diet-disturbed gut microbiota-colonocyte interactions contribute to dysregulating peripheral tryptophan-kynurenine metabolism. Microbiome 11, 154 (2023). 10.1186/s40168-023-01606-x

76 Matsumoto, M., Kakizoe, K. & Benno, Y. Comparison of fecal microbiota and polyamine concentration in adult patients with intractable atopic dermatitis and healthy adults. Microbiol Immunol 51, 37–46 (2007). 10.1111/j.1348-0421.2007.tb03888.x

77 Matsumoto, M. et al. Impact of intestinal microbiota on intestinal luminal metabolome. Sci Rep 2, 233 (2012). 10.1038/srep00233

78 Matsumoto, M. & Benno, Y. The relationship between microbiota and polyamine concentration in the human intestine: a pilot study. Microbiol Immunol 51, 25–35 (2007). 10.1111/j.1348-0421.2007.tb03887.x

79 Nakamura, A., Ooga, T. & Matsumoto, M. Intestinal luminal putrescine is produced by collective biosynthetic pathways of the commensal microbiome. Gut Microbes 10, 159–171 (2019). 10.1080/19490976.2018.1494466

80 Lux, G. D., Marton, L. J. & Baylin, S. B. Ornithine decarboxylase is important in intestinal mucosal maturation and recovery from injury in rats. Science 210, 195–198 (1980). 10.1126/science.6774420

81 Timmons, J., Chang, E. T., Wang, J. Y. & Rao, J. N. Polyamines and Gut Mucosal Homeostasis. J Gastrointest Dig Syst 2 (2012).

82 Nakamura, A. et al. Symbiotic polyamine metabolism regulates epithelial proliferation and macrophage differentiation in the colon. Nat Commun 12, 2105 (2021). 10.1038/s41467-021-22212-1

83 Puleston, D. J. et al. Polyamines and eIF5A Hypusination Modulate Mitochondrial Respiration and Macrophage Activation. Cell Metab 30, 352–363 e358 (2019). 10.1016/j.cmet.2019.05.003

84 Hardbower, D. M. et al. Ornithine decarboxylase regulates M1 macrophage activation and mucosal inflammation via histone modifications. Proc Natl Acad Sci U S A 114, E751–E760 (2017). 10.1073/pnas.1614958114

85 Bardocz, S., Grant, G., Brown, D. S. & Pusztai, A. Putrescine as a source of instant energy in the small intestine of the rat. Gut 42, 24–28 (1998). 10.1136/gut.42.1.24

86 Grancara, S. et al. Spermine cycling in mitochondria is mediated by adenine nucleotide translocase activity: mechanism and pathophysiological implications. Amino Acids 48, 2327–2337 (2016). 10.1007/s00726-016-2264-6

87 Jing, Y. H. et al. Spermidine ameliorates the neuronal aging by improving the mitochondrial function in vitro. Exp Gerontol 108, 77–86 (2018). 10.1016/j.exger.2018.04.005

88 Phillips, J. E. & Chaffee, R. R. Restorative effects of spermine on oxidative phosphorylation and respiration in heat-aged mitochondria. Biochem Biophys Res Commun 108, 174–181 (1982). 10.1016/0006-291x(82)91847-2

89 Nicchitta, C. V. & Williamson, J. R. Spermine. A regulator of mitochondrial calcium cycling. J Biol Chem 259, 12978–12983 (1984).

90 Pezzato, E., Battaglia, V., Brunati, A. M., Agostinelli, E. & Toninello, A. Ca2+-independent effects of spermine on pyruvate dehydrogenase complex activity in energized rat liver mitochondria incubated in the absence of exogenous Ca2+ and Mg2+. Amino Acids 36, 449–456 (2009). 10.1007/s00726-008-0099-5

91 Levy, M. et al. Microbiota-Modulated Metabolites Shape the Intestinal Microenvironment by Regulating NLRP6 Inflammasome Signaling. Cell 163, 1428–1443 (2015). 10.1016/j.cell.2015.10.048

92 Ahmadi, S. et al. A human-origin probiotic cocktail ameliorates aging-related leaky gut and inflammation via modulating the microbiota/taurine/tight junction axis. JCI Insight 5 (2020). 10.1172/jci.insight.132055

93 Holecek, M. Histidine in Health and Disease: Metabolism, Physiological Importance, and Use as a Supplement. Nutrients 12 (2020). 10.3390/nu12030848

94 Barcik, W. et al. Histamine-secreting microbes are increased in the gut of adult asthma patients. J Allergy Clin Immunol 138, 1491–1494 e1497 (2016). 10.1016/j.jaci.2016.05.049

95 Duraski, R. M. Ciprofloxacin-induced theophylline toxicity. South Med J 81, 1206 (1988). 10.1097/00007611-198809000-00042

96 Zhao, Y. et al. Histamine Intolerance-A Kind of Pseudoallergic Reaction. Biomolecules 12 (2022). 10.3390/biom12030454

97 Schnedl, W. J. & Enko, D. Histamine Intolerance Originates in the Gut. Nutrients 13 (2021). 10.3390/nu13041262

98 Schink, M. et al. Microbial patterns in patients with histamine intolerance. J Physiol Pharmacol 69 (2018). 10.26402/jpp.2018.4.09

99 Sanchez-Perez, S. et al. Intestinal Dysbiosis in Patients with Histamine Intolerance. Nutrients 14 (2022). 10.3390/nu14091774

100 Raithel, M., Matek, M., Baenkler, H. W., Jorde, W. & Hahn, E. G. Mucosal histamine content and histamine secretion in Crohn’s disease, ulcerative colitis and allergic enteropathy. Int Arch Allergy Immunol 108, 127–133 (1995). 10.1159/000237129

101 Winterkamp, S. et al. Urinary excretion of N-methylhistamine as a marker of disease activity in inflammatory bowel disease. Am J Gastroenterol 97, 3071–3077 (2002). 10.1111/j.1572-0241.2002.07028.x

102 Hagel, A. F. et al. Plasma histamine and tumour necrosis factor-alpha levels in Crohn’s disease and ulcerative colitis at various stages of disease. J Physiol Pharmacol 66, 549–556 (2015).

103 Smolinska, S. et al. Histamine Receptor 2 is Required to Suppress Innate Immune Responses to Bacterial Ligands in Patients with Inflammatory Bowel Disease. Inflamm Bowel Dis 22, 1575–1586 (2016). 10.1097/MIB.0000000000000825

104 Maintz, L. & Novak, N. Histamine and histamine intolerance. Am J Clin Nutr 85, 1185–1196 (2007). 10.1093/ajcn/85.5.1185

105 Schwelberger, H. G., Hittmair, A. & Kohlwein, S. D. Analysis of tissue and subcellular localization of mammalian diamine oxidase by confocal laser scanning fluorescence microscopy. Inflamm Res 47 **Suppl 1**, S60–61 (1998). 10.1007/s000110050273

106 Petersen, J., Raithel, M. & Schwelberger, H. G. Histamine N-methyltransferase and diamine oxidase gene polymorphisms in patients with inflammatory and neoplastic intestinal diseases. Inflamm Res 51 **Suppl 1**, S91–92 (2002). 10.1007/pl00022464

107 Schulze, H. A. et al. From model cell line to in vivo gene expression: disease-related intestinal gene expression in IBD. Genes Immun 9, 240–248 (2008). 10.1038/gene.2008.11

108 Kojima, S. & Ichibagase, H. Studies on synthetic sweetening agents. 8. Cyclohexylamine, a metabolite of sodium cyclamate. Chem Pharm Bull (Tokyo) 14, 971–974 (1966). 10.1248/cpb.14.971

109 Reuter, S. E. & Evans, A. M. Carnitine and acylcarnitines: pharmacokinetic, pharmacological and clinical aspects. Clin Pharmacokinet 51, 553–572 (2012). 10.1007/BF03261931

110 Evans, A. M. & Fornasini, G. Pharmacokinetics of L-carnitine. Clin Pharmacokinet 42, 941–967 (2003). 10.2165/00003088-200342110-00002

111 Knottnerus, S. J. G. et al. Disorders of mitochondrial long-chain fatty acid oxidation and the carnitine shuttle. Rev Endocr Metab Disord 19, 93–106 (2018). 10.1007/s11154-018-9448-1

112 El-Gharbawy, A. & Vockley, J. Inborn Errors of Metabolism with Myopathy: Defects of Fatty Acid Oxidation and the Carnitine Shuttle System. Pediatr Clin North Am 65, 317–335 (2018). 10.1016/j.pcl.2017.11.006

113 Roediger, W. E. The colonic epithelium in ulcerative colitis: an energy-deficiency disease? Lancet 2, 712–715 (1980). 10.1016/s0140-6736(80)91934-0

114 Roediger, W. E. & Nance, S. Metabolic induction of experimental ulcerative colitis by inhibition of fatty acid oxidation. Br J Exp Pathol 67, 773–782 (1986).

115 Peltekova, V. D. et al. Functional variants of OCTN cation transporter genes are associated with Crohn disease. Nat Genet 36, 471–475 (2004). 10.1038/ng1339

116 Smith, S. A. et al. Mitochondrial dysfunction in inflammatory bowel disease alters intestinal epithelial metabolism of hepatic acylcarnitines. J Clin Invest 131 (2021). 10.1172/JCI133371

117 Barrett, J. C. et al. Genome-wide association defines more than 30 distinct susceptibility loci for Crohn’s disease. Nat Genet 40, 955–962 (2008). 10.1038/ng.175

118 Singh, R. et al. Autophagy regulates lipid metabolism. Nature 458, 1131–1135 (2009). 10.1038/nature07976

119 Lemons, J. M. S. et al. Enterobacteriaceae Growth Promotion by Intestinal Acylcarnitines, a Biomarker of Dysbiosis in Inflammatory Bowel Disease. Cell Mol Gastroenterol Hepatol 17, 131–148 (2024). 10.1016/j.jcmgh.2023.09.005

120 Balasubramanian, K. et al. Metabolism of the colonic mucosa in patients with inflammatory bowel diseases: an in vitro proton magnetic resonance spectroscopy study. Magn Reson Imaging 27, 79–86 (2009). 10.1016/j.mri.2008.05.014

121 Ooi, M. et al. GC/MS-based profiling of amino acids and TCA cycle-related molecules in ulcerative colitis. Inflamm Res 60, 831–840 (2011). 10.1007/s00011-011-0340-7

122 Schicho, R. et al. Quantitative metabolomic profiling of serum, plasma, and urine by (1)H NMR spectroscopy discriminates between patients with inflammatory bowel disease and healthy individuals. J Proteome Res 11, 3344–3357 (2012). 10.1021/pr300139q

123 Stephens, N. S. et al. Urinary NMR metabolomic profiles discriminate inflammatory bowel disease from healthy. J Crohns Colitis 7, e42–48 (2013). 10.1016/j.crohns.2012.04.019

124 Alonso, A. et al. Urine metabolome profiling of immune-mediated inflammatory diseases. BMC Med 14, 133 (2016). 10.1186/s12916-016-0681-8

